# Differences in gene expression in field populations of *Wolbachia*-infected *Aedes aegypti* mosquitoes with varying release histories in northern Australia

**DOI:** 10.1101/2022.10.10.511644

**Authors:** B.M.C. Randika Wimalasiri-Yapa, Bixing Huang, Perran A. Ross, Ary A. Hoffmann, Scott A. Ritchie, Francesca D. Frentiu, David Warrilow, Andrew F. van den Hurk

**Affiliations:** Department of Medical Laboratory Sciences, Faculty of Health Sciences, Open University of Sri Lanka, Nugegoda, Colombo, Sri Lanka; School of Biomedical Sciences and Centre for Immunology and Infection Control, Queensland University of Technology, Brisbane, Queensland, Australia; Public Health Virology, Forensic and Scientific Services, Department of Health, Queensland Government, Coopers Plains, Queensland, Australia; Pest and Environmental Adaptation Research Group, School of BioSciences, Bio21 Institute, The University of Melbourne, Melbourne, Victoria, Australia; Australian Institute of Tropical Health and Medicine, James Cook University, Cairns, Queensland, Australia

## Abstract

*Aedes aegypti* is the principal mosquito vector of dengue, yellow fever, Zika and chikungunya viruses. The *w*Mel endosymbiotic bacteria *Wolbachia pipientis* has been introduced into this vector as a novel biocontrol strategy to stop transmission of these viruses. Mosquitoes with *Wolbachia* have been released in the field in North Queensland, Australia since 2011, at various locations and over several years, with populations remaining stably infected. *Wolbachia* infection is known to alter gene expression in its mosquito host, but whether (and how) this changes over the long-term in the context of field releases remains unknown. We sampled mosquitoes from *Wolbachia*-infected populations with different release histories along a time gradient and performed RNAseq to investigate gene expression changes in the insect host. We observed a significant impact on gene expression in *Wolbachia*-infected mosquitoes versus uninfected controls, but fewer genes had altered expression in the older releases (e.g. the year 2011) versus the more recent releases (e.g. 2017). Nonetheless, a fundamental signature of *Wolbachia* infection on host gene expression was observed across all releases, comprising upregulation of immunity and metabolism genes. There was limited downregulation of gene expression in the older releases, but significantly more in the most recent release. Our findings indicate that at > 8 years post-introgression into field populations, *Wolbachia* continues to profoundly impact host gene expression, particularly genes involved in insect immune response. We suggest that if gene expression changes underlie blocking of virus replication in *Wolbachia*-infected *Ae. aegypti*, then refractoriness of these mosquitoes to arboviruses may remain stable over the long-term.

**Author summary:** The *Aedes aegypti* mosquito is the main species responsible for urban transmission of dengue, Zika and chikungunya viruses. Control measures, including source reduction and insecticide treatment, have historically struggled to provide sustained control of this species to limit disease. An alternative approach involves releasing mosquitoes harbouring *Wolbachia* bacteria. *Wolbachia* inhibits virus transmission by *Ae. aegypti* and preliminary evidence indicates that dengue incidence is reduced in locations where it has been deployed. In this study, we found that *Wolbachia* significantly upregulates gene expression in *Ae. aegypti* at least 8 years after field deployment compared with uninfected controls, although some gene downregulation was also observed. We observed a more ‘muted’ response in mosquitoes from populations with older release histories, with far fewer genes being differentially regulated versus those from the most recent release. Irrespective of release history, immune response and metabolism genes were significantly upregulated, and to a lesser extent genes related to behaviour. Our results, combined with previous studies that have revealed few changes in the *Wolbachia* genome post release, provide further evidence of the long-term stability of the *Wolbachia*-mosquito relationship in the field.

## Introduction

The *Aedes aegypti* mosquito is the primary urban vector of dengue, yellow fever, chikungunya and Zika viruses [1]. *Aedes aegypti* has a close association with humans, who are its primary blood meal source [2]. It also preferentially blood-feeds and rests indoors, and utilizes water-filled receptacles proximal to human habitation as larval habitats. These behaviours make *Ae. aegypti* notoriously difficult to control, particularly in high-density urban areas [3]. Control of *Ae. aegypti* has relied on the reduction, elimination or insecticide treatment of receptacle habitats and/or application of adulticides, either as space sprays or targeted indoor residual application [4]. However, lack of sustainability of current government-administered control programs, coupled with insecticide resistance, compromises effectiveness of these control strategies. More sustainable approaches to controlling *Ae. aegypti* and its associated arboviruses are required, especially due to a lack of suitable vaccines (except for yellow fever) or antiviral therapies to limit disease.

Alternative strategies involving the targeted release of modified *Ae. aegypti* are at various stages of development and show excellent promise for sustained control of arbovirus transmission [5, 6]. One of the most advanced involves the release of *Ae. aegypti* transinfected with the endosymbiotic bacterium *Wolbachia pipientis*, which confers phenotypes that can be exploited to limit arbovirus transmission [7]. High rates of maternal transmission and cytoplasmic compatibility (CI) allow the bacteria to spread through and be maintained in the resident mosquito population [8, 9]. Arbovirus replication and transmission is blocked in mosquitoes infected with *Wolbachia* [10-12]. Following initial success with driving the *w*Mel strain of *Wolbachia* into *Ae. aegypti* populations in urban centers of north Queensland, Australia [8, 13, 14], releases of *Wolbachia*-infected *Ae. aegypti* are being conducted in at least 12 countries [9, 15]. Epidemiological evidence suggests a significant reduction in dengue incidence post deployment in dengue endemic locations [9, 16].

Stability of the endosymbiont infection in *Ae. aegypti* will be critical for the long-term viability of *Wolbachia*-based arbovirus control programs. Any loss of the virus blocking phenotype could lead to increased virus transmission by mosquitoes. High temperatures could lead to the loss of *Wolbachia* from populations [17-19], but other factors (such as high fitness costs due to *Wolbachia* or the release strain and dry environmental conditions) could be responsible too [20, 21]. Alternatively, there are three potential vulnerabilities for breakdown of virus blocking related to the evolution of microbe and mosquito: virus evolutionary escape, and changes to the *Wolbachia* or mosquito genomes. Hence, post-release long-term monitoring should include periodic genome sequencing and assessment of the virus, bacteria and mosquito. As dengue viruses have an RNA genome, they are subject to relatively high mutational rates compared to other DNA-based organisms and microbes, so selection of virus strains which escape from the effects of *Wolbachia* are a possibility [22].

Other evolutionary pressures may drive genetic changes in either the *Wolbachia* or the mosquito. In recent studies to explore the first possibility, mosquitoes collected from north Queensland had few changes in their *Wolbachia* genome sequences compared to the pre-release strain, indicating a high level of stability to date [23, 24]. In addition, a comparison of the mosquito genomes also suggests that there have been few changes in the mosquito genome since *Wolbachia* replacement [25], whilst the frequency of *Wolbachia* in invaded populations has remained high [14, 26] and most host effects of the Wolbachia have remained stable [26] with the exception of effects on egg quiescence [27]. Although few genomic changes have so far been detected and linked to structural changes in genes, it is possible that there have nevertheless been changes in the expression of host genes which are often sensitive indicators of adaptation, including in immune responses [28].

Changes in gene expression patterns of *Wolbachia*-infected mosquitoes could lead to loss of virus blocking abilities. Hypotheses on a virus blocking mechanism can be grouped into the two broad categories of mosquito immune gene activation and/or competition for host cell resources (reviewed in [29, 30]. However, immune genes are activated by transinfected *Wolbachia* but are not required for blocking in naturally infected hosts, so may not be essential [31, 32]. Important examples of antiviral pathways in the insect cell include the activation of signalling pathways such as Janus kinase-signal transducer and activator of transcription (JAK-STAT), reactive oxygen species (ROS), and Toll signalling. Various anti-microbial proteins and compounds can be induced such as the interferon-like Vago and Dnmt2 which may modify viral RNA, making it susceptible to methylation-mediated degradation. However, none of these pathways has so far been directly linked to the blocking effect [30]. The exonuclease XRN1 is induced in *Wolbachia*-infected cells and has been associated with viral RNA degradation [33]. RNA interference is also activated but may not be an important viral blocking pathway in *Wolbachia*-infected cells [34].

An alternative hypothesis to immune activation is that virus is in competition with *Wolbachia* for physical space and other host cell resources. *Wolbachia* density in some cases may be correlated with the virus blocking effect [35, 36], and exclusion of virus from cells and tissues where *Wolbachia* is found at high density suggests competition for cellular resources. Cell nutrients are regulated by the host, and *Wolbachia* [37] and viruses may compete for amino acids. *Wolbachia* may also alter the metabolism of lipids such as cholesterol [38, 39], a molecule which is also essential for dengue virus replication [40]. Alternatively, *Wolbachia* may physically exclude RNA viruses from cellular organelles necessary for viral replication [41]. The relative contributions of immune activation and host resource competition in virus blocking remains an open question. Elucidating whether immune activation persists in long-term *Ae. aegypti* - *Wolbachia* associations would shed light on this question and inform considerations of the long-term stability of virus blocking.

The objectives of the study described here were threefold. First, we obtained samples from multiple locations in and around Cairns, Australia, the site of releases of *Wolbachia*-infected mosquitoes from 2011 – 2017 and compared their gene expression to uninfected mosquitoes to determine which genes were differentially expressed. Second, we established a baseline for gene expression in *Wolbachia*-infected mosquitoes for longer-term studies which will help monitor for any breakdown in virus blocking. Finally, *Wolbachia*-infected mosquitoes were present at some sites for up to eight years, so we determined if there were any trends or differences in mosquito gene expression between sites with older compared with more recent releases.

## Results

### Mosquito gene expression clusters according to *Wolbachia* infection status

Eggs were collected in 2019 from populations where *Wolbachia*-infected *Ae. aegypti* were released in years 2011 (Aae.wMel_2011_), 2013-2014 (Aae.wMel_2013/2014_) and 2017 (Aae.wMel_2017_), as well as from a wild-type population (Aae.wt) where *Wolbachia*-infected mosquitoes had not been released (Fig 1). Eggs were hatched and reared under laboratory conditions and transcriptome sequencing (RNA-Seq) of pools of 5 adult females at 4 days post emergence was performed (Table 1).

**Table 1.** *Aedes aegypti* collected in April 2019 from Cairns, Queensland, Australia, for mosquito transcriptome sequencing.

| Suburb | Release date<sup>a</sup> | No. sample sites | No. mosquitoes | No. pools |
| --- | --- | --- | --- | --- |
| Gordonvale | January 2011 | 10 | 50 | 10 |
| Yorkeys Knob | January 2011 | 2 | 10 | 2 |
| Edge Hill | January 2013 | 8 | 40 | 8 |
| Parramatta Park | January 2013 | 1 | 5 | 1 |
| Cairns North | August 2014 | 1 | 5 | 1 |
| Bungalow | July 2014 | 1 | 5 | 1 |
| Cairns North | March 2017 | 3 | 15 | 3 |
| Parramatta Park | March 2017 | 2 | 10 | 2 |
| Caravonica | No release | 5 | 25 | 5 |

<sup>a</sup>Date when releases of *Wolbachia*-infected *Ae. aegypti* commenced in the suburb.

**Figure 1.**
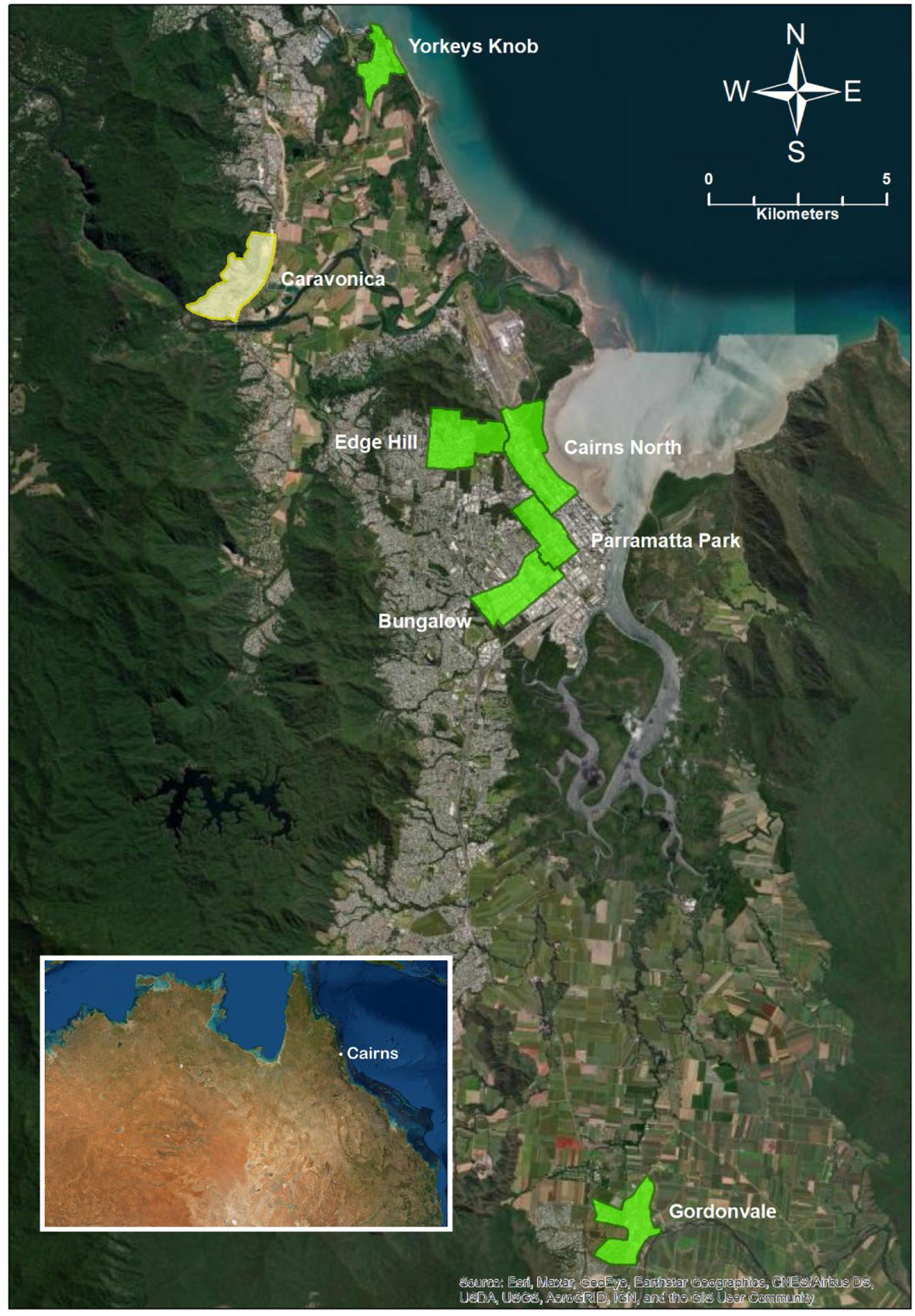
Map of mosquito collection locations in the Cairns Regional Council area, northern Queensland, Australia. *Wolbachia*-infected mosquitoes were released in the suburbs shaded green. There had been no releases of *Wolbachia* infected mosquitoes in Caravonica (shaded yellow) up until at least April 2019, when the collections for the current study were undertaken.

To ensure that the variation in gene expression described below was likely due to actual gene expression and not differences in *Wolbachia* density, the number of copies of *w*Mel *wsp* gene in *Ae. aegypti* was quantified using the method of Lee et al. [42]. There was no significant difference (Mann-Whitney test *P* = 0.5926) in *Wolbachia* density between mosquitoes collected from the 2013-14 and 2017 release locations (S1 Fig; note that there was no material available for density quantification from the 2011 release sites).

A total of 2,041,416,107 Illumina raw reads were obtained from sequencing on the Novaseq at the Australian Genome Research Facility (AGRF). Among these, there was a total of 1,554,852,005 read pairs and overall 89.42% of the pairs were aligned to the *Ae. aegypti* reference genome GCF_002204515.2 (https://www.ncbi.nlm.nih.gov/genome/?term=txid7159[orgn]) (see S1 Table for mapping statistics). Following initial processing of raw reads (see Methods), differentially expressed genes (DEGs) were identified by comparing gene expression in *Wolbachia*-infected versus uninfected mosquitoes using the Deseq2 tool. DEGs were considered statistically significant when the absolute fold change of the gene expression was > 2 and adjusted *P*-value was < 0.05. Comparison of gene expression between all infected mosquitoes (n = 28 pools) and uninfected mosquitoes (n = 5 pools) resulted in a total of 747 DEGs (656 upregulated and 91 downregulated). A heatmap of differential expression in these 747 genes indicated separate clustering of *Wolbachia*-infected and uninfected mosquitoes (Fig 2).

**Figure 2.**
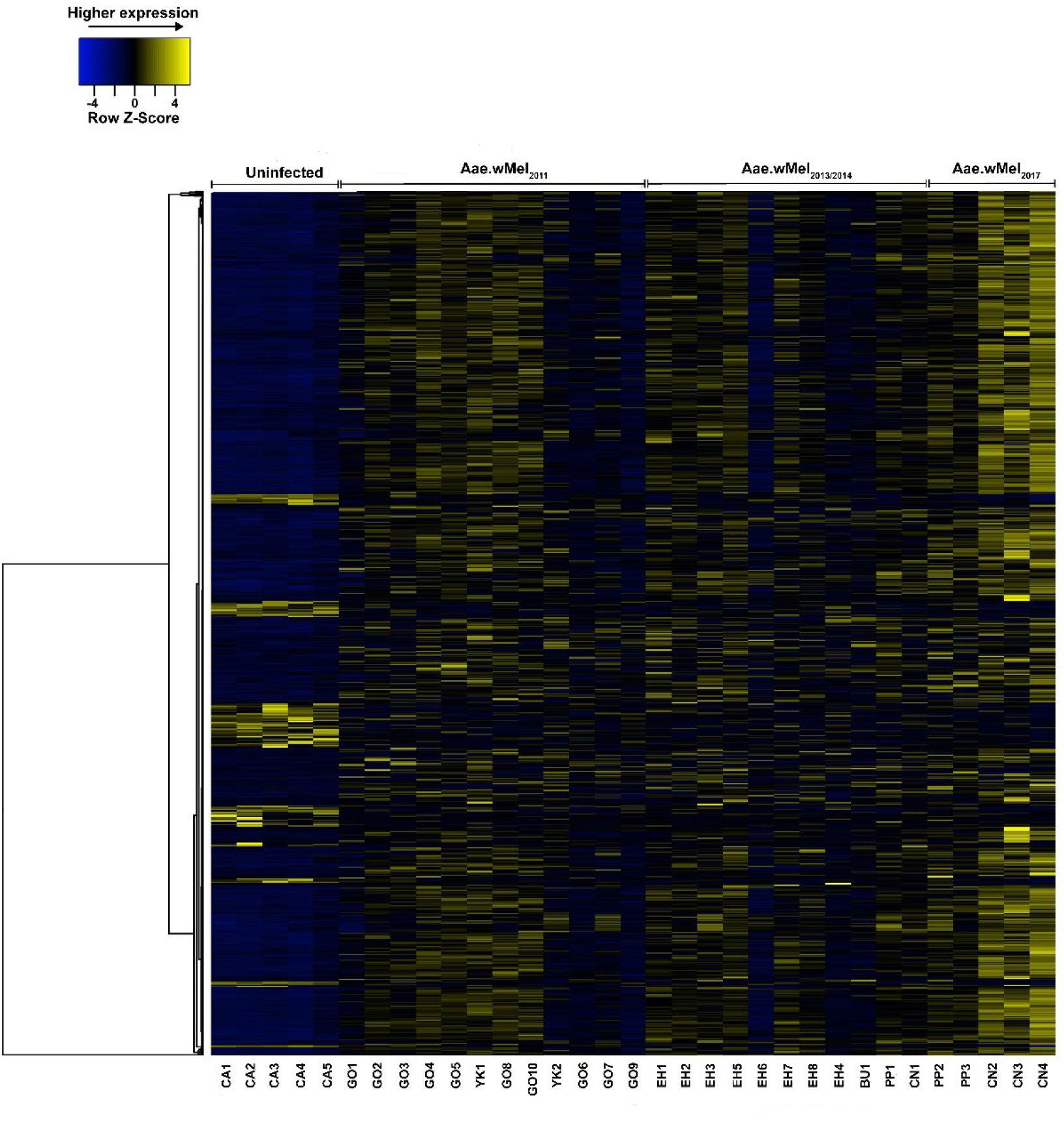
Heatmap of normalised read counts of differentially expressed genes (DEGs) showing clustering of transcriptomic response in *w*Mel *Wolbachia*-infected *Aedes aegypti* from locations with different release years versus uninfected *Ae. aegypti*. Hierarchical clustering was performed using the complete linkage method and the distances between columns were computed by the Euclidean method (http://www.heatmapper.ca/expression/). Acronyms for the individual mosquito pool identifiers on the X-axis denote suburb names (CA = Caravonica, GO = Gordonvale, YK = Yorkeys Knob, EH = Edge Hill, BU = Bungalow, PP = Parramatta Park, and CN = Cairns North).

### Elevated DEGs in mosquitoes from the most recent *Wolbachia* field release

An analysis of the gene expression of *Wolbachi*a-infected versus uninfected *Ae. aegypti* indicated that the highest number of DEGs was observed in the Aae.wMel_2017_ mosquitoes (i.e. mosquitoes descended from releases that occurred 2 years prior to collection for this study (Table 2)). By contrast, a smaller number of DEGs was observed in mosquitoes from older releases, Aae.wMel_2011_ and Aae.wMel_2013/2014_ (Table 2), relative to *Wolbachia*-uninfected mosquitoes. The difference in number of DEGs from the Aae.wMel_2017_ mosquitoes versus those from the Aae.wMel_2011_ and Aae.wMel_2013/2014_ mosquitoes was statistically significant (Chi-square *P* < 0.001). A principal component analysis (PCA) of all genes indicated a clear demarcation of gene expression between infected and uninfected mosquitoes descended from 2017 releases (Fig 3). This demarcation was less apparent in mosquitoes with earlier release histories.

**Table 2.**
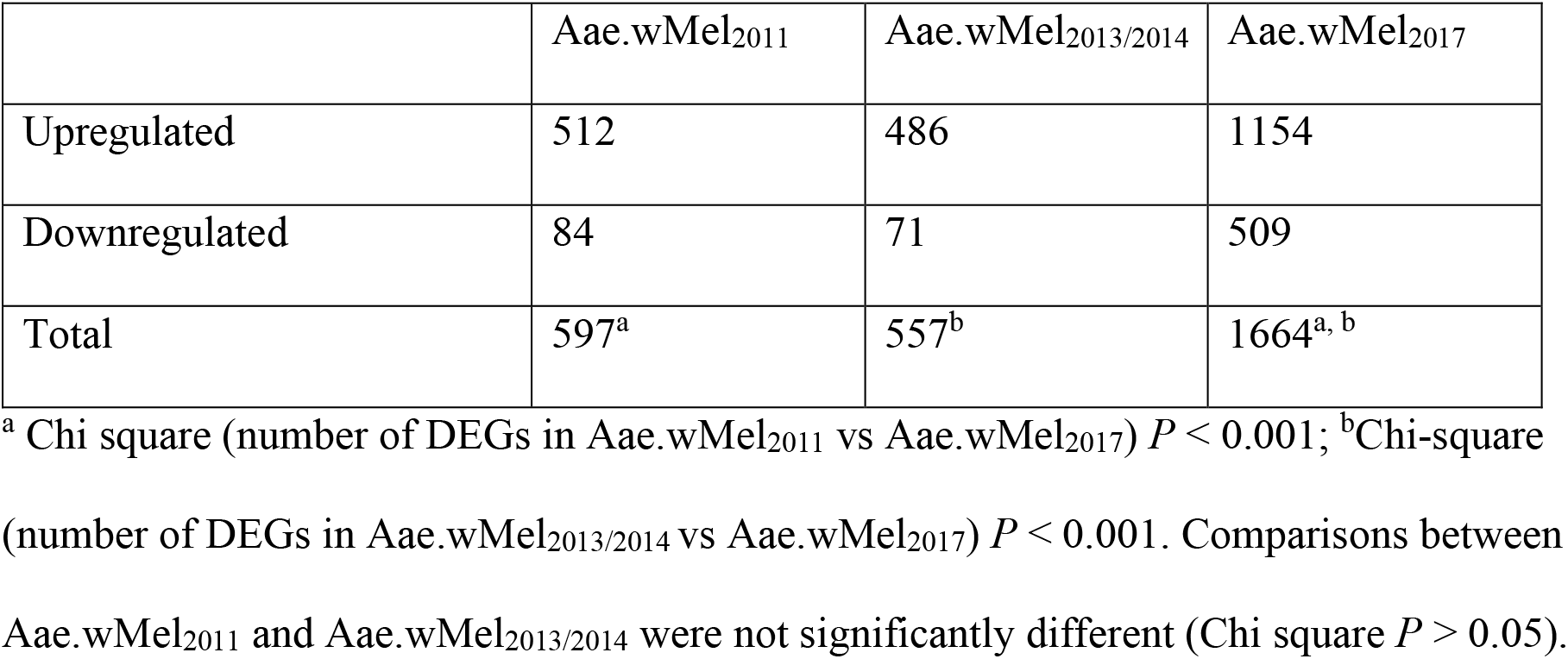
Number of DEGs (absolute fold change ±2 and adjusted *P*-value (false discovery rate (FDR)) ≤ 0.05) in mosquitoes from populations with different *w*Mel release years versus *Wolbachia*-uninfected mosquitoes.

|  | Aae.wMel<sub>2011</sub> | Aae.wMel<sub>2013/2014</sub> | Aae.wMel<sub>2017</sub> |
| --- | --- | --- | --- |
| Upregulated | 512 | 486 | 1154 |
| Downregulated | 84 | 71 | 509 |
| Total | 597<sup>a</sup> | 557<sup>b</sup> | 1664<sup>a, b</sup> |

<sup>a</sup> Chi square (number of DEGs in Aae.wMel<sub>2011</sub> vs Aae.wMel<sub>2017</sub>) $P < 0.001$ ; <sup>b</sup> Chi-square (number of DEGs in Aae.wMel<sub>2013/2014</sub> vs Aae.wMel<sub>2017</sub>) $P < 0.001$ . Comparisons between Aae.wMel<sub>2011</sub> and Aae.wMel<sub>2013/2014</sub> were not significantly different (Chi square $P > 0.05$ ).

**Figure 3.**
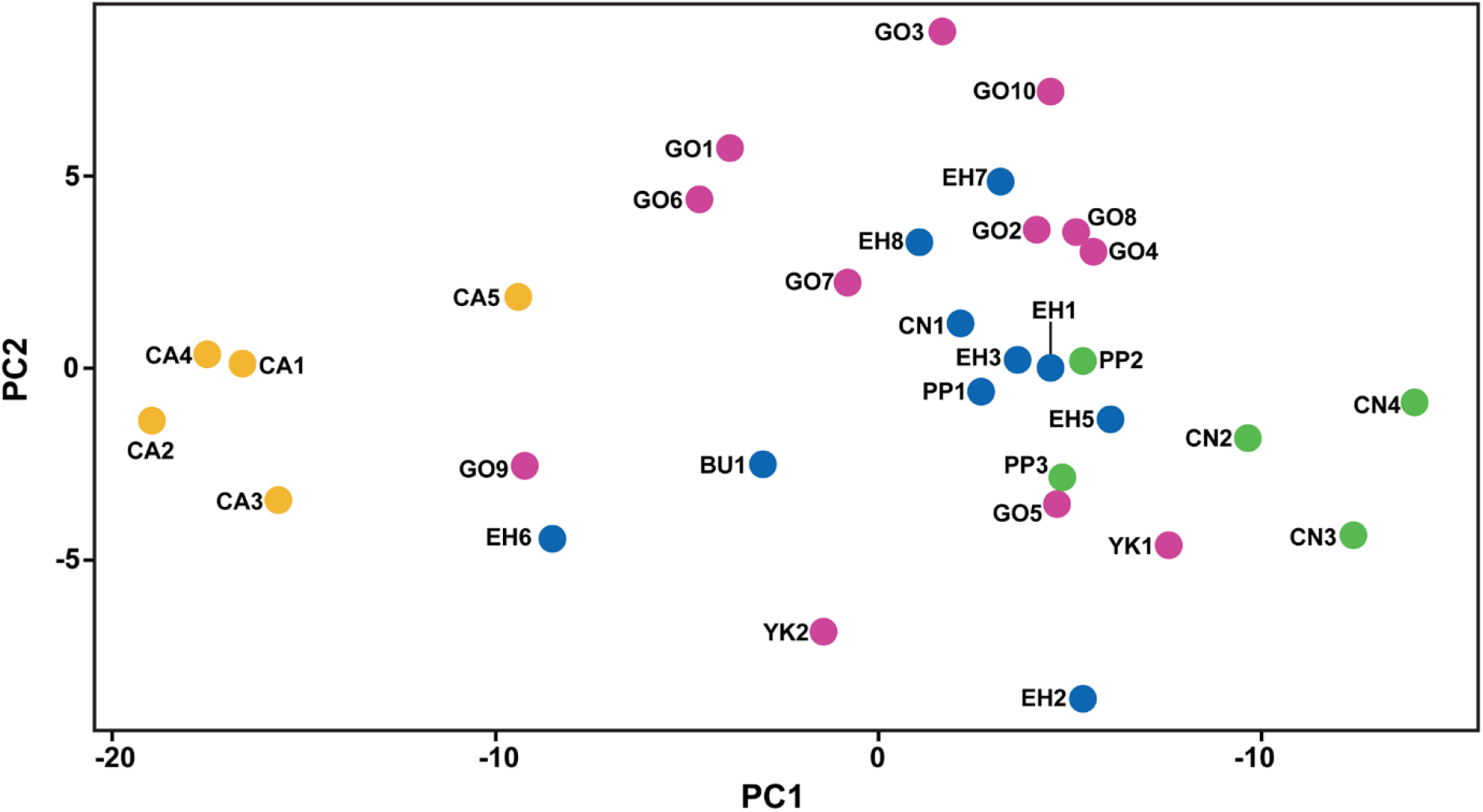
Principal component analysis (PCA) of gene expression in *Wolbachia*-infected mosquitoes originating from different release histories: Aae.wMel_2017_ vs uninfected (green); Aae.wMel_2013/2014_ vs uninfected (blue); and Aae.wMel_2011_ vs uninfected (pink). Uninfected mosquitoes (yellow shading) were from the suburb of Caravonica where releases of *Wolbachia*-infected mosquitoes had not been conducted up until the time of our collections in April 2019. Acronyms for the individual mosquito pool identifiers denote suburb names (CA = Caravonica, GO = Gordonvale, YK = Yorkeys Knob, EH = Edge Hill, BU = Bungalow, PP = Parramatta Park, and CN = Cairns North).

Comparison of DEGs from the three *Wolbachia*-infected mosquito release times identified 357 common upregulated genes and 23 common downregulated genes (Fig 4). There were 76, 37 and 1029 unique DEGs for the mosquitoes from *w*Mel releases in the Aae.wMel_2011_, Aae.wMel_2013/2014_ and Aae.wMel_2017_ mosquitoes, respectively, compared with the uninfected mosquitoes.

**Figure 4.**
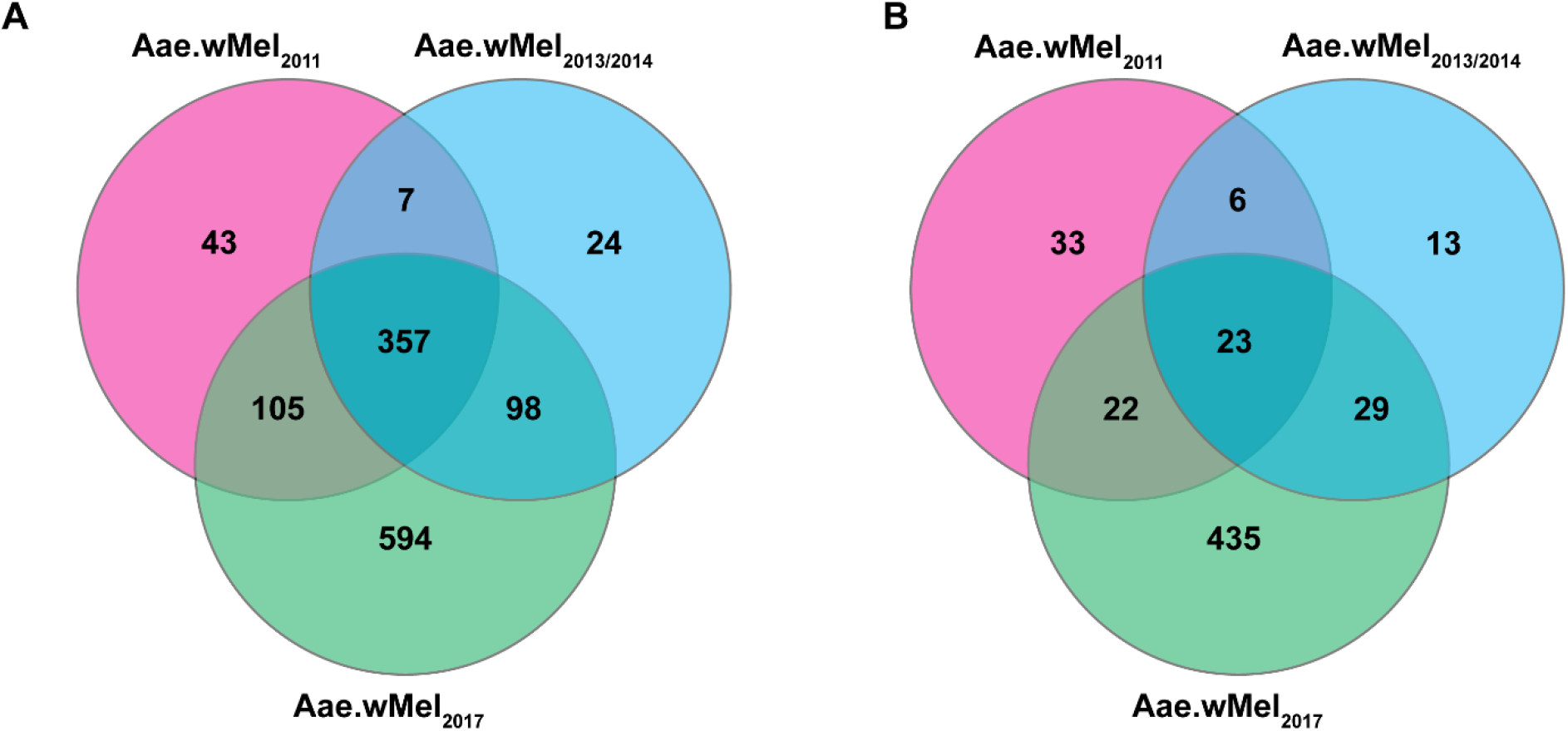
Numbers of shared and uniquely (A) upregulated and (B) downregulated genes in mosquitoes from populations with different *w*Mel release histories.

We identified 594 genes to be uniquely upregulated in the mosquitoes from locations of *w*Mel releases in 2017 (that is, not found in any of the earlier releases). The comparison of expression level of these 594 genes across all populations using the Kruskal-Wallis test showed that expression of these genes was significantly associated with time point (the year that mosquito population acquired *w*Mel) (*P* < 0.001). Post-hoc analysis revealed that the median gene expression (counts per million) in Aae.wMel_2017_ mosquitoes was significantly higher than those originating from the earlier releases (*P* < 0.001; Table 3).

**Table 3.** Pairwise comparison of median gene expression (CPM) according to release history.

| Pairwise comparison of CPM | Mean difference | *P*-value* |
| --- | --- | --- |
| Aae.wMel<sub>2017</sub> vs Aae.wMel<sub>2013/2014</sub> | 9035.44 vs 8161.84 | <0.001 |
| Aae.wMel<sub>2013/2014</sub> vs Aae.wMel<sub>2011</sub> | 8161.84 vs 8158.72 | 0.970 |
| Aae.wMel<sub>2017</sub> vs Aae.wMel<sub>2011</sub> | 9035.44 vs 8158.72 | <0.001 |

\**P* value based on post-hoc analysis of Kruskal-Wallis test.

### Upregulated DEGs with the highest fold change belong to immune response

Immune function genes, genes associated with stress response and non-coding genes comprised the 10 most upregulated genes, with the highest fold change values in comparisons involving *Wolbachia*-infected from all release populations versus uninfected mosquitoes. Among these, the majority (70-80%, depending on release history) were immune genes (Fig 5). Three immune genes were significantly upregulated in all *w*Mel-infected mosquito populations, irrespective of release year. These were CTLGA8 (LOC5575053), alpha-2-macroglobulin (LOC23687443) and leucine-rich repeat-containing protein 40-like (LOC110677030). There were 7, 8 and 8 DEGs related to immune response in mosquitoes from Aae.wMel_2011_, Aae.wMel_2013/2014_ and Aae.wMel_2017_, respectively. The topmost upregulated gene in Aae.wMel_2011_ and Aae.wMel_2013/2014_ mosquitoes was the transferrin gene (LOC5579417), with this gene being the second-most upregulated in Aae.wMel_2017_ mosquitoes. Two out of three remaining topmost upregulated genes of Aae.wMel_2011_ mosquitoes were CLIP (LOC5578693) and leucine-rich repeat (LOC5575814), both pathogen recognition receptor (PRR) molecules that are important for immune function. The other was arrestin C-terminal-like domain-containing protein 3 (LOC5570224) which plays a role in regulation of the olfactory system (33). Similarly, Aae.wMel_2013/2014_ and Aae.wMel_2017_ mosquitoes also show two PRRs each including LOC5575053 and LOC110677006 in the earlier, and LOC5570871 and LOC5576315 in the later release. Additionally, mosquitoes from these releases showed differential expression of putative defense protein 1 (LOC5572918) and a negative regulator of translation (LOC5569955). Altogether, four genes (2011: LOC5567033, LOC5566857; 2013-14: LOC110673980; and 2017: LOC5563952) out of the 10 topmost upregulated genes across all three mosquito groups are regarded as involved in stress response while two genes were non-coding (Fig 5).

**Figure 5.**
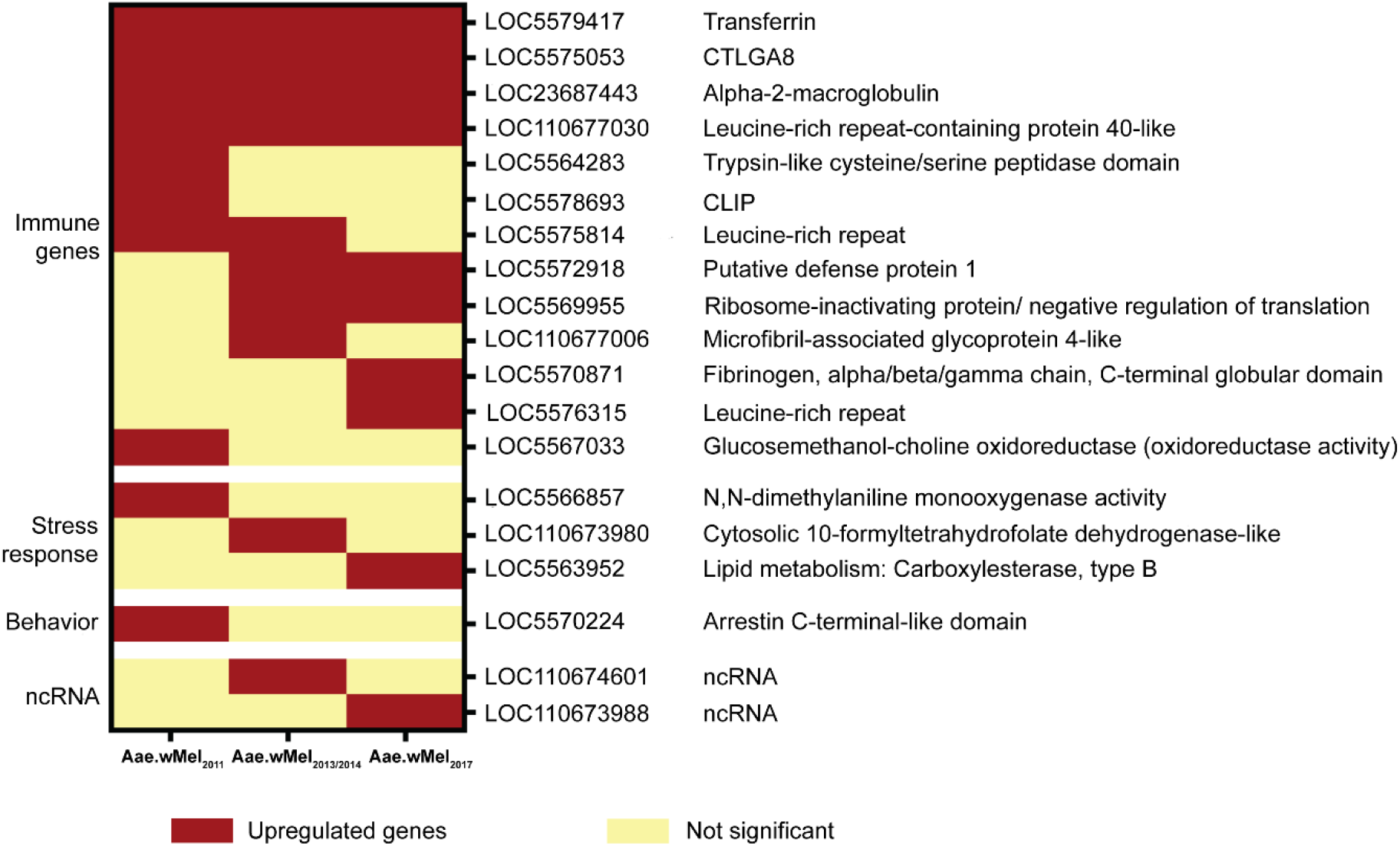
The top 10 upregulated genes with highest fold difference between *w*Mel infected and uninfected *Ae. aegypti*, according to release year. The type of upregulated gene (immune genes, stress response, ncRNA, and behaviour) is shown.

### Topmost downregulated genes belong to non-coding RNAs, cell proliferation and host behaviour

The highest fold downregulated gene categories included non-coding RNAs, genes involved in cell replication, and host behaviour related genes (Fig 6). Unlike upregulated genes with highest fold changes mentioned above, none of the downregulated genes were common to mosquitoes from all three *Wolbachia* release histories. However, four downregulated DEGs common to Aae.wMel_2011_ and Aae.wMel_2013/2014_ mosquitoes were CFI06_mgr02, LOC5574234, LOC5576517 and LOC5572259. Aae.wMel_2011_ had the majority (6/10) of downregulated genes with the highest fold change in either non-coding RNA genes (LOC110676610, LOC110679144, LOC5574600, LOC110676459) or uncharacterized genes (LOC5572259, LOC5568345). Three out of 10 topmost downregulated genes with highest fold change from Aae.wMel_2013/2014_ were either ncRNA or uncharacterized proteins. Genes that were downregulated with highest fold change in Aae.wMel_2017_ mosquitoes were unique except for histone-lysine N-methyltransferase Suv4-20 (LOC5569935) which was shared with Aae.wMel_2013/2014_ mosquitoes. Some of these genes included three ncRNA genes (LOC110679860, LOC5564187, LOC5563860), a neurotransmitter gene (LOC23687658), a gene responsible for promotion of micropinocytosis (LOC110674232) and three genes playing a role in cell replication.

**Figure 6.**
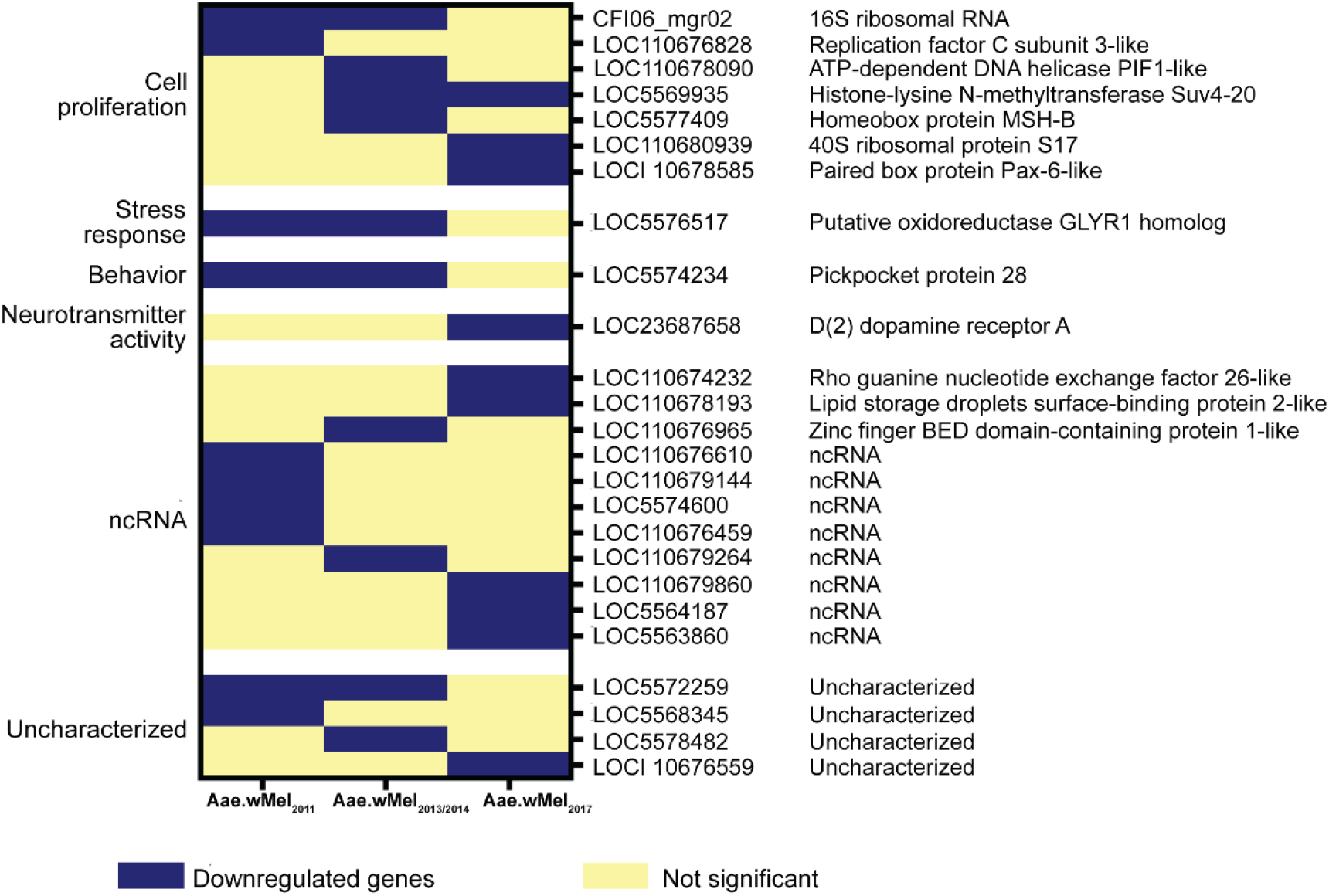
Topmost downregulated genes with highest fold difference between *w*Mel infected and uninfected *Ae. aegypti*, according to release year. The type of downregulated gene (cell proliferation, stress response, behaviour, neurotransmitter activity, ncRNA, and uncharacterised) is shown.

### *w*Mel is associated with upregulation of pathways related to immunity, amino acid and lipid metabolism, and behaviour, across all mosquito release time points

Gene ontology (GO) analysis performed using the DAVID bioinformatics tool [43] revealed that 357 upregulated genes common to all releases resulted in six significantly enriched (Fisher’s exact *P* <0.05) biological processes including innate immune response (GO:0045087), fatty acid biosynthetic process (GO:0006633), phototransduction (GO:0007602), defense response to bacterium (GO:0042742), visual perception (GO:0007601) and urea cycle (GO:0000050) (Fig 7). Moreover, two cellular locations comprising extracellular space (GO:0005615) and extracellular region (GO:0005576) were significantly enriched at the same cut-off criterion. Molecular functions, namely serine-type endopeptidase activity (GO:0004252) and endopeptidase inhibitor activity (GO:0004866) were significantly enriched (Fisher’s exact *P* <0.05).

**Figure 7.**
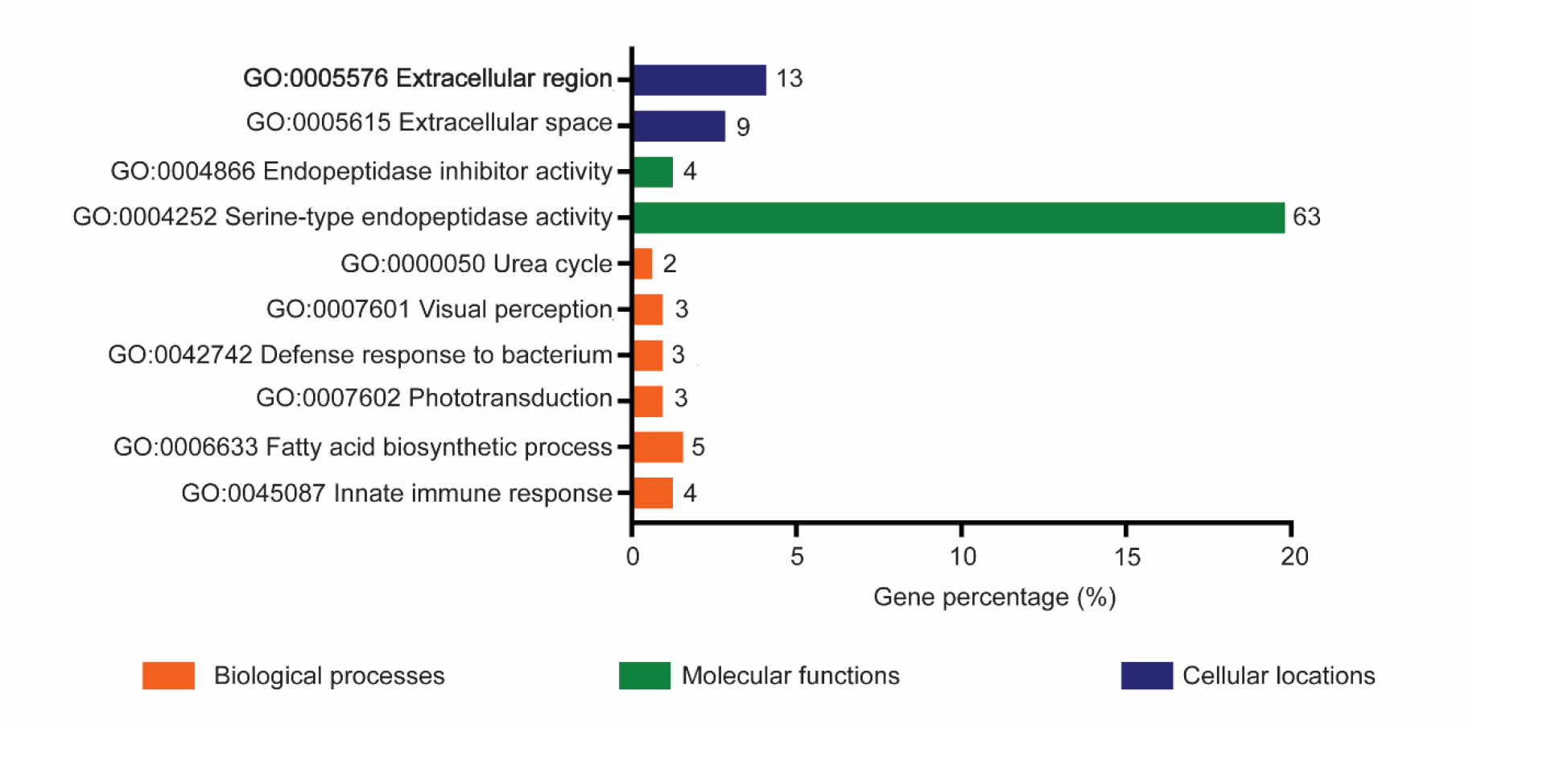
Gene ontology analysis of upregulated differentially expressed genes common to all *Wolbachia*-infected *Ae. aegypti* versus uninfected mosquitoes. The number next to each bar refers to the number of genes involved in the biological process, molecular function or cellular location.

Kyoto Encyclopedia of Genes and Genomes (KEGG) pathway enrichment analysis (FDR<0.05) of these proteins further revealed that metabolic pathways (aag01100), glycine, serine and threonine metabolism (aag00260), biosynthesis of amino acids (aag01230), arginine biosynthesis (aag00220), tyrosine metabolism (aag00350), arginine and proline metabolism (aag00330), caffeine metabolism (aag00232), carbon metabolism (aag01200) and phototransduction – fly (aag04745) were upregulated (see S2 Table for enrichment analysis data).

### Uniquely upregulated DEGs in Aae.wMel2017 mosquitoes are involved in stress response, membrane transport and iron metabolism

Following the observation of higher levels of DEGs in the Aae.wMel_2017_ mosquitoes versus other release histories, we explored what types of genes were present in this group, starting with upregulated genes. For this purpose, first we evaluated upregulated DEGs (n=594) unique to Aae.wMel_2017_ mosquitoes. Three GO terms of biological process, namely sodium ion transport (GO:0006814), ion transport (GO:0006811) and metabolic process (GO:0008152), were significantly enriched at Fisher’s exact *P* < 0.05. Moreover, four GO terms of cellular locations: integral component of membrane (GO:0016021), extracellular region (GO:0005576), gap junction (GO:0005921) and extracellular space (GO:0005615) were also significantly enriched. Cytochrome-c oxidase activity (GO:0004129), oxidoreductase activity (GO:0016491), monooxygenase activity (GO:0004497), oxidoreductase activity, acting on paired donors, with incorporation or reduction of molecular oxygen (GO:0016705), iron ion binding (GO:0005506), serine-type endopeptidase activity (GO:0004252), hemebinding (GO:0020037), transporter activity (GO:0005215), phosphopantetheine binding (GO:0031177) and hydrolase activity (GO:0016787) were among the significantly enriched molecular functions.

KEGG enrichment (FDR < 0.05) indicated several pathways that were significantly enriched in Aae.wMel_2017_ mosquitoes, but not the other release years. Starch and sucrose metabolism (aag00500), other glycan degradation (aag00511), glyoxylate and dicarboxylate metabolism (aag00630), glycine, serine, and threonine metabolism (aag00260), oxidative phosphorylation (aag00190), glycolysis / gluconeogenesis (aag00010), galactose metabolism (aag00052), amino sugar and nucleotide sugar metabolism (aag00520) and lysosome (aag04142) were among the uniquely enriched pathways. Moreover, additional genes of KEGG pathways such as biosynthesis of amino acids (aag01230), carbon metabolism (aag01200) and metabolic pathways (aag01100) were also identified during KEGG pathway enrichment analysis.

### Downregulated DEGs in Aae.wMel2011 mosquitoes are related to DNA replication

Eighty-four downregulated DEGs were uploaded for GO analysis using the DAVID bioinformatics tool. Nucleosome assembly was the only GO term pertaining to biological processes that was significantly enriched (Fisher’s exact *P* <0.05). Two GO terms, Nucleus (GO:0005634) and nucleosome (GO:0000786), were the significantly enriched cellular locations at Fisher’s exact *P* <0.05. None of the GO term classified as molecular functions (GO:0003677: DNA binding) was significantly enriched (Fisher exact *P* > 0.05). Three functional annotation clusters resulted from a DAVID cluster analysis, in which the first cluster had three GO terms (GO:0005634: nucleus, GO:0000786: nucleosome and GO:0003677: DNA binding), four UNIPROT keywords (DNA-binding, nucleosome core, chromosome, and nucleus) and an InterPro term (IPR009072: histone-fold). The second cluster included one GO term (GO:0008270: zinc ion binding) and two InterPro terms (IPR011011: Zinc finger, FYVE/PHD-type and IPR013083: Zinc finger, RING/FYVE/PHD-type). The third cluster was included with three UniProt keywords: transmembrane helix, transmembrane and membrane; one GO term: GO:0016021: integral component of membrane.

A considerable number (36/84) of genes were unmapped to the DAVID cloud map gene IDs. Among these unmapped genes, 16 were non-coding RNA genes (S3 Table) and three were uncharacterized, while information for the rest is given in the S4 Table.

### Downregulated DEGs in Aae.wMel2013/2014 and Aae.wMel2017 mosquitoes are related to multicellular organism development

When considering Aae.wMel_2013/2014,_ there were 71 downregulated DEGs, among which 26 genes (36.6%) of the Gene IDs were not identified in the DAVID cloud map. There was a single biological process significantly enriched at Fisher’s exact *P* < 0.05: multicellular organismal process (GO:0032501). None of the cellular locations or molecular processes were significantly enriched. DAVID pathway enrichment analysis identified four KEGG pathways that were enriched: notch signalling pathway (aag04330), lysine degradation (aag00310), dorso-ventral axis formation (aag04320) and Wnt signalling pathway (aag04310). However, none of these were enriched at Fisher’s *P* < 0.05. There were three annotation clusters that were enriched at a score > 0.5 in DAVID functional annotation clustering. The first cluster included three UniProt keywords (Receptor, Transducer and G-protein coupled receptor) and two InterPro protein families (IPR000276: G protein-coupled receptor, rhodopsin-like, IPR017452: GPCR, rhodopsin-like, 7TM). The second cluster included one GO term (GO:0016021 integral component of membrane) and three UniProt key words (transmembrane helix, transmembrane and membrane). Two UniProt keywords (DNA-binding and nucleus) and one GO term (GO:0005634 nucleus) comprised the third cluster. Genes that were not mapped to DAVID cloud map were manually checked using an NCBI Gene search (S5 Table). Eleven out of 26 were ncRNA (S3 Table). Three genes were uncharacterised (LOC110674591, LOC110674325 and LOC110680306), while six out of remaining 12 genes were transcription factors (LOC110675182, LOC110678581, LOC110675146, LOC110678585, LOC110676930 and LOC110674313).

Among 509 DEGs observed in Aae.wMel_2017_ mosquitoes, 405 downregulated DEGs were significantly enriched for 12 biological processes and 10 molecular functions in DAVID bioinformatics analysis (S1 Box). Moreover, cellular locations such as nucleus (GO:0005634), nucleosome (GO:0000786), MCM complex (GO:0042555) and origin recognition complex (GO:0000808) were significantly enriched at Fisher’s exact *P* < 0.05. Genes that were not mapped to DAVID cloud map were manually checked using an NCBI Gene search (S6 Table). There were 104 downregulated DEGs that were not mapped to the DAVID cloud map, among which 34 were ncRNA (S3 Table) and 11 were uncharacterised (LOC110676333, LOC110678663, LOC110674783, LOC110678024, LOC110681172, LOC110680396, LOC110681054, LOC110681485, LOC110676281, LOC110676629, LOC110675455 and LOC110679465). The other DEGs were basically related to immune response, cell proliferation and development (S6 Table)

## Discussion

Our transcriptome analysis has identified DEGs in *w*Mel-infected *Ae. aegypti* descended from mosquitoes released in Cairns, Australia, in 2011, 2013-14 and 2017. We found that there was a significantly higher number of DEGs in Aae.wMel_2017_ mosquitoes compared with Aae.wMel_2011_ and Aae.wMel_2013/2014_ mosquitoes. There are several potential explanations for the differences between years. The results of the quantitative PCR analysis suggest that it is likely not associated with overall *Wolbachia* density within mosquitoes. It is plausible that gene expression in subsets of genes has become attenuated as the mosquito undergoes evolutionary changes in response to *Wolbachia* infection [25]. Previous studies have demonstrated the attenuation of *Wolbachia*-mediated phenotypes (particularly CI, fecundity and fitness effects) in *Drosophila* spp. infected with the “popcorn” strain of *Wolbachia* [44, 45] although most traits associated with *w*Mel appear to be phenotypically stable [26].

The DEGs that we have identified may give insight into the nature of the virus-blocking response, as well as other phenotypes that *Wolbachia* confers to transinfected mosquitoes. If the virus blocking phenotype is underpinned by the expression of a gene, or set of genes, rather than structural modifications to host cells by *Wolbachia*, this is more likely to be differentially expressed in all release groups (i.e. the common DEGs) than are other genes. Importantly then, common DEGs may reveal clues to identity and function of the virus blocking gene(s). Hence, our analysis has focussed on characterising the DEGs common to all the release groups. However, there is a caveat that virus blocking genes may not be expressed in mosquitoes descended from all release years as much as other DEGs for the following reasons: the RNA-Seq methodology may not detect all differentially expressed genes, sampling may be insufficient, their temporal expression may differ, or expression may vary such that a difference cannot be detected. Hence, some analysis of the type and function of DEGs not common to all the release groups is warranted but is only discussed briefly below. Finally, as Aae.wMel_2017_ mosquitoes had significantly higher numbers of both up- and down-regulated DEGs compared with the other release years, the gene expression changes in that group will also be discussed to help understand this observation which may be related to ongoing host evolution of *Wolbachia* mediated effects [27].

Broadly, DEGs common to all release years were categorized into immune responses, metabolic changes, and cell proliferation gene-function groups. Innate immune priming is one possible mechanism of viral blocking by *Wolbachia* infected mosquitoes [32], with protection provided to the insect by pre-activation or upregulation of antimicrobial encoding genes [30]. There are four types of genes that are involved in immune responses in mosquitoes: pattern recognition receptors (PRRs), activation of immune signalling, immune effector mechanisms and immune modulation by the regulation of mosquito homeostasis [46]. In this study, the genes responsible for pathogen recognition, such as CTLGA8, alpha-2-macroglobulin and leucine-rich repeat-containing protein 40-like were among the top 10 differentially expressed genes with highest fold change. In addition, among the common DEGs found in all *w*Mel-infected mosquitoes analysed here, there were numerous types of insect PRRs identified, including leucine rich repeats, galectins, CLIP domain serine proteases, scavenger receptor proteins, peptidoglycan recognition protein (short), gram negative bacteria binding proteins and C-type lectins. Similar findings were previously identified in a study on gene expression in *Anopheles gambiae* cells during *Wolbachia* infection [47].

Previous studies have indicated that *Wolbachia* induces the Toll pathway via reactive oxygen species to mediate oxidative stress and activate antimicrobial peptides, defensins and cecropins, and other Toll pathway genes [10, 32]. In our study, we also identified antimicrobial peptides such as cecropin E, defensin-C and defensin-D and diptericin 1, two PRRs such as gram-negative bacteria-binding protein 1(GNBP1), peptidoglycan-recognition protein 2 (PGRP1) and one other gene (uncharacterized protein LOC5577955 and isoform X1) that are related to Toll and IMD pathway to be significantly upregulated in all *w*Mel-infected *Ae. aegypti*. In this study, the iron binding protein, transferrin 1 which is suggested to have functions in iron metabolism and immune function [48], was the upregulated DEG with either highest or second highest fold change in the mosquitoes from all release histories. Transferrin-1 gene upregulation in response to *Wolbachia* infection in mosquitoes has been previously reported [32, 49], while iron dependence of *Wolbachia* on different host species has also been identified [50-52]. This iron sequestration from the host has also been suggested as a pathogen blocking mechanism via alteration of iron binding during DENV and Zika infection of *Ae. aegypti* [53].

The upregulation of lysozyme genes that degrade pathogens was also evident in our study. It has previously been shown that DENV infection triggers autophagy which initiates lysosomal degradation of the virus [54]. Autophagy is a conserved mechanism that degrades cellular components to maintain tissue homeostasis [55], and has been reported in other dipterans. For instance, antiviral activation of autophagy in *Drosophila* spp. has been reported against vesicular stomatitis virus and Rift Valley fever virus [56, 57]. In our study, we also identified a cathepsin gene and the KEGG pathway lysosome (aag04142) that were upregulated in *w*Mel-infected *Ae. aegypti* compared with uninfected mosquitoes, and these are potentially involved in autophagy [58]. Our study also provides evidence of stability of pre-activation of autophagy response in *Wolbachia*-infected mosquitoes at least up to 8 years after *w*Mel is introgressed into wild populations. Altogether, our study indicates that many immune components including PRRs, signalling pathways and immune effectors are significantly altered in gene expression in mosquitoes infected with *w*Mel, and this response is conserved at least 8 years after initial invasion of the endosymbiont into natural populations.

Competition for host cellular resources has been suggested as another mechanism of virus blocking induced by *Wolbachia* infection [30, 37, 38], as *Wolbachia* is dependent on the mosquito cell for lipid and amino acid biosynthesis [29]. In our study, we also observed the upregulation of genes related to fatty acid metabolism including fatty acid synthase, elongation of very long chain fatty acids protein 4 and 7, and acyl-CoA Delta (11) desaturase isoform X2 as well as myeloid differentiation 2-related lipid recognition protein. When considering DEGs of amino acid metabolism, our study revealed that all *w*Mel-infected *Ae. aegypti* had significantly upregulated glycine, serine and threonine metabolism (aag00260), biosynthesis of amino acids (aag01230), arginine biosynthesis (aag00220), tyrosine metabolism (aag00350), arginine and proline metabolism (aag00330). It has been previously demonstrated that cysteine, glutamate, glutamine, proline, serine and threonine are used as energy sources by *w*Mel [59], and *Ae. aegypti* fecundity and egg viability was affected by competition with *Wolbachia* for amino acids [37]. In another study, leucine, tryptophan, methionine, valine, histidine, lysine, phenylalanine, arginine, asparagine and threonine were found to be essential for successful egg production while cysteine, glycine and isoleucine were considered semi-essential for egg production [60]. Overall, upregulation of the genes related to amino acid metabolism in all *w*Mel-infected *Ae. aegypti* in our study supports previous studies which have shown the effects of *Wolbachia* infection on insect host physiology and metabolism, particularly in *Ae. aegypti* transinfected with the *w*MelPop strain [37]. *Ae. aegypti* does not encode a functional urea cycle [61], with mosquitoes regulating urea metabolism through an interaction between the argininolysis and uricolysis pathways [62, 63]. Our study provides evidence of upregulation of genes related to the argininolysis pathway in all *Ae. aegypti* infected with *w*Mel, while the involvement of uricolysis pathway is evident in Aae.wMel_2017_ mosquitoes. It is possible that enhanced metabolic requirements due to *Wolbachia* presence may lead to a greater need for nitrogen excretion, but this requires further study.

Previous studies have identified that myeloid differentiation 2-related lipid recognition protein (ML) and the Niemann Pick-type C1 (NPC1), which are lipid binding proteins, act as agonists of DENV infection by altering vesicular trafficking, lipid metabolism and the endoplasmic reticulum to facilitate viral entry and replication [64, 65]. Specifically, clathrin-mediated endocytosis was identified to be the entry pathway of DENV into mosquito cells [66-68]. KEGG Brite hierarchical clustering of DEGs upregulated in all *w*Mel infected mosquitoes in our study indicated that some genes responsible for membrane trafficking were upregulated. Those genes were low density lipoprotein receptor adapter protein 1-A which is involved in clathrin-mediated endocytosis, perlucin-like protein a C-lectin receptor which is involved in phagocytosis, and glutamyl aminopeptidase, an endoplasmic reticulum (ER)-Golgi intermediate compartment (ERGIC) protein which is involved in ER-Golgi transport forward pathways. Therefore, our study suggests dysregulation of membrane proteins could be a contributor to membrane-mediated virus blocking but the mechanism remains to be elucidated.

Our analysis identified that some downregulated genes were related to cell proliferation including transcription and translation and, additionally, organism development. For instance, Aae.wMel_2011_ mosquitoes showed downregulation of genes involved in head or eye development, development of the clypeolabrum and several head sensory organs (protein optix-like: LOC110680308) [69, 70], nerve cord development (protein commissureless 2 homolog: LOC 110679455) [71] and interneuron differentiation (zinc finger C4H2 domain-containing protein-like: LOC110678839) [72]. DEGs downregulated in Aae.wMel_2013/2014_ included paired box protein Pax-6-like gene (LOC110678585) which is involved in eye morphogenesis [73], ATP-dependent DNA helicase PIF1-like genes (LOC110678090) which are DNA replication proteins (https://www.genome.jp/kegg-bin/get_htext?aag00001+5580192), and mantle protein-like (LOC110676076), which had 98% similarity to Vajk2 gene by protein Blast, which plays a role in chitin-based cuticle development in *Drosophila melanogaster* [74]. In addition to these, when we included the downregulated DEGs in KEGG mapper analysis, we identified that there were some commonly affected pathways in the Aae.wMel_2013/2014_ and Aae.wMel_2017_ mosquitoes. Those included Notch signalling, Wnt signalling and mTOR pathways, in addition to pheromone/odorant binding protein genes which were commonly identified in all *w*Mel infected mosquitoes. Notably, there was a significantly higher number of genes downregulated in Aae.wMel_2017_ than Aae.wMel_2011_ and Aae.wMel_2013/2014_ mosquitoes. Specifically, Aae.wMel_2017_ mosquitoes showed downregulation of pathways that control cell replication. These observations may be tied to *Wolbachia*’s reliance on the host cell, for example its manipulation of the cytoskeleton, to achieve successful replication [29]. A previous study has also observed the involvement of cell replication pathways and Notch signalling in *w*Mel-infected mosquitoes when subjected to selection on dengue virus-blocking [75].

We also found the E3 SUMO-protein ligase protein inhibitor of activated STAT (PIAS) 2-like (LOC110674232) gene to be downregulated. Interestingly, a previous study identified that silencing of this gene makes *Ae. aegypti* more resistant to DENV-2 [76] as PIAS is the negative regulator of the JAK-STAT pathway [77]. Forkhead-box (Fox) transcription factors are found to play an important role in reproduction of *Ae. aegypti* [78]. Specifically, the knockdown of the FoxO gene affected the activation of amino acid mediated expression of vitellogenin gene and resulted in a decrease in egg deposition/egg numbers by female *Ae. aegypti*. The above-mentioned study further suggested the possibility of regulation of the amino acid target of rapamycin (TOR) pathway via Fox genes including FoxO. The mosquito mTOR pathway is found to play a key role in regulation of complete vitellogenesis [79, 80] whilst Wnt signalling is identified as the main event in *Drosophila* development [81].

It was noted that several non-coding RNAs (ncRNAs) appeared among the top-most up- and down-regulated DEGs. It is increasingly appreciated that long ncRNAs are important in various biological processes including but not limited to cell differentiation, epigenetic and non-epigenetic based gene regulation, involvement in the defence system, responses to stimuli and stress response, viral replication and antiviral defence [82-88]. Our study also identified ncRNA loc110674601, which is significantly aligned with Arginine-glutamic acid dipeptide repeats protein (blastn alignment not shown). This protein plays a role as a transcriptional repressor during mouse development and in the control of cell survival [89]. Loc110673988 (AAEL022454) was among the top 10 upregulated DEG in Aae.wMel2017. A previous study has indicated that this gene is involved in mosquito cellular immunity [90]. There were eight ncRNA among the topmost downregulated genes. Importantly, the downregulated genes with highest fold change in Aae.wMel_2011_ (LOC110676610) and Aae.wMel_2017_ (LOC110679860: ncRNA) were ncRNAs. Our data suggest that ncRNAs may play hitherto unappreciated roles in the ability of *Wolbachia* to successfully colonise the mosquito host.

It important to note that whilst every effort was made to sample as many locations as possible in our study, only approximately half of the traps contained sufficient numbers of *Ae. aegypti* to include in our analysis. This limited our ability to compare gene expression differences between mosquitoes in geographically separate suburbs for a given release date. Furthermore, although we show different gene expression profiles between different release dates, future sampling should be conducted in other global release locations, such as Yogyakarta [16] or Kuala Lumpur [9], to examine whether geographically separate local release sites show geographical and temporal differences in gene expression. Changes in gene expression related to other processes, such as the shift in quiescence patterns [27], also require further study.

## Conclusions

There is a general decrease in the number of DEGs as a result of *w*Mel *Wolbachia* infection with time post-release. However, *w*Mel infection is characterized by a prolonged transcriptomic signature with respect to upregulated genes (up to 8 years) while downregulated gene signatures were partially fixed until 5-6 years. Upregulated genes and pathways in the host associated with *w*Mel infection were mainly related to immunity and metabolism (especially amino acid and lipid metabolism), while downregulated genes were related to odorant/pheromone binding and reproduction/ organism development. Our results suggest that *w*Mel-based vector control method is robust for at least 8 years with respect to immune priming and host resources-depletion based pathogen blocking, if these are indeed the mechanisms responsible for this phenomenon. This fixed gene signature comprises transcriptomic alterations in immunity, stress response, behavior and metabolic changes. There were also effects on genes associated with host reproduction which were strongest in the most recent releases but not evident after 8 years.

## Materials and methods

### Sample collection, RNA extraction and cDNA library preparation

*Aedes aegypti* eggs were collected from the Cairns suburbs of Caravonica, Gordonvale, Yorkeys Knob, Edge Hill, Parramatta Park, Bungalow and Cairns North in April 2019 (Fig 1 and Table 1). At the time of sampling, Caravonica was one of the few remaining locations with *Wolbachia*-free *Ae. aegypti*.

Field-collected eggs were reared under standard insectary conditions as described by Huang et al. [24]. Post emergence, adult mosquitoes were maintained on 15% honey water as a nutrient source. On day 4 post emergence, they were anaesthetised on wet ice and sorted by species and sex. Females were washed in absolute EtOH before being placed in RNALater (Qiagen) and stored at -80ºC.

RNA was extracted from pools of 5 females using an RNeasy Mini kit (Qiagen) by first homogenizing them using a plastic pestle in 600 µL of lysis buffer in a 1.5 mL microfuge tube. RNA was then extracted following the manufacturer’s recommended method and the presence of *Wolbachia* in the pool tested using previously outlined methods [24]. The RNA was analysed on a TapeStation and quality assessed by determination of the RNA integrity number (RIN). Samples with RIN scores less than 7.8 were excluded from further analysis. A polyadenylated fraction was purified from the total RNA (1 µg) using the NEBNext® Poly(A) mRNA Magnetic Isolation Module (New England Biolabs). This fraction was used to construct cDNA library using a previously described method [91]. Briefly, poly(A) RNA (2-5 ng) was converted to cDNA using the Protoscript II kit (New England Biolabs) and a supplied mix of random hexamer and d(T)_23_VN primers, followed by conversion to double-stranded cDNA using a cocktail of RNase H, DNA ligase and DNA polymerase I (New England Biolabs). The product was used to construct a barcoded cDNA library using the Nextera XT system (Illumina) which was sequenced on NovaSeq 6000 at the Australian Genome Research Facility (AGRF), generating paired 2x 150 nt reads. A total of approximately 60 million reads were obtained for each sample. Reads are available from the NCBI Short Read Archive under Accession Number PRJNA867516 (a private link is provided here for review purposes only: https://dataview.ncbi.nlm.nih.gov/object/PRJNA867516?reviewer=cn61lep54dl2j0bqdi2qug89tp).

### Bioinformatic analysis

Raw transcriptomic data (616.51Gb) were uploaded to Galaxy [92] Australia cloud and subjected to quality control using fastqc tool [93]. Reads from four lanes were merged using concatenate tail-to-head (cat) as per R1 and R2 and then trimmed and adaptor sequences were removed. Reads with quality Phread score < 30 and read length < 50 were excluded using the Trim Galore tool. Next, the *Ae. aegypti* reference genome GCF_002204515.2 was downloaded from NCBI and mapped to the trimmed sequence pairs using Hisat2. Gene expression was quantified using the feature counts tool. Differentially expressed genes were then identified using Deseq2 (FDR< 0.05 and absolute fold change ± 2) by comparing gene expression of *w*Mel *Ae. aegypti* released at different times (2011, 2013-14 and 2017) against wild type *Ae. aegypti*. Downstream analysis of upregulated genes was performed after comparing gene lists using Venny 2.1.0-BioinfoGP. Gene ontology (GO) analysis was performed using the DAVID bioinformatics tool, with GO terms identified to be significantly enriched using Fisher’s exact *P*-value < 0.05 [43].

Upregulated DEGs that were common to all *w*Mel *Ae. aegypti* populations were input into KEGG mapper (https://www.genome.jp/kegg/tool/map_pathway2.html) to identify altered gene expression pathways. Downstream analysis of downregulated genes was performed according to the mosquito population with DAVID bioinformatics tool. Any DEG that was not identified by the DAVID bioinformatics cloud map characterised manually by searching either NCBI or VectorBase gene search, or protein/nucleotide blast.

Normality of gene expression values were assessed by the Shapiro-Wilks test, as implemented in SPSS software (IBM). As expected, gene expression values (counts per million-CPM) were not normally distributed. Thus, the non-parametric Kruskal-Wallis test was used to check relationships between gene expression and timepoint of release, as implemented in SPSS using a *P*-value of < 0.05 to determine statistical significance.

### Quantification of *Wolbachia* in *Ae. aegypti*

To ensure that the differences observed above were due to gene regulation to attenuate a costly immune and/or metabolic detoxoxification response and not due to changes in *Wolbachia* density, the *w*Mel in *Ae. aegypti* collected from 2013-14 and 2017 release locations was quantified using the quantitative PCR developed by Lee et al. [42]. The *Wolbachia* density was analysed using a Mann-Whitney U test in GraphPad Prism Version 9.1.0 [94].

## Acknowledgements

We wish to thank Tom Schmidt (University of Melbourne) and Michael Townsend (Australian Institute of Tropical Health and Medicine, James Cook University) for collecting the mosquito eggs used in this study.

## Supporting information

**S1 Fig. Density of *Wolbachia* in *Aedes aegypti* descended from mosquitoes released in the Cairns region of northern Australia in 2013-14 and 2017**. Each dot is an individual mosquito, and bars and whiskers are medians and 95% confidence intervals, respectively. There was no significant difference (*P* > 0.05; Mann-Whitney U test) in *Wolbachia* density between the years.

**S1 Table. Mapping statistics**.

**S2 Table. KEGG pathway enrichment analysis of 357 commonly upregulated DEGs across all time points**.

**S3 Table. Unmapped downregulated DEGs which were non-coding RNA from *Aedes aegypti* with different release years**.

**S4 Table. Unmapped downregulated DEGs in Aae.wMel**_**2011**_ **mosquitoes which are not ncRNA**.

**S5 Table. Unmapped downregulated DEGs from Aae.wMel**_**2013/2014**_ **mosquitoes.**

**S1 Box. Significantly enriched GO terms pertaining to biological processes and molecular functions in Aae.wMel**_**2017**_ **mosquitoes**.

**S6 Table. Unmapped downregulated DEGs in Aae.wMel**_**2017**_ **mosquitoes**.

