## Supplemental files for "Differences in gene expression in field populations of *Wolbachia*-infected *Aedes aegypti* mosquitoes with varying release histories in northern Australia"

### Supporting information

**S1 Fig. Density of *Wolbachia* in *Aedes aegypti* descended from mosquitoes released in the Cairns region of northern Australia in 2013-14 and 2017.** Each dot is an individual mosquito and bars and whiskers are medians and 95% confidence intervals, respectively. There was no significant difference ($P > 0.05$; Mann-Whitney U test) in *Wolbachia* density between the years.

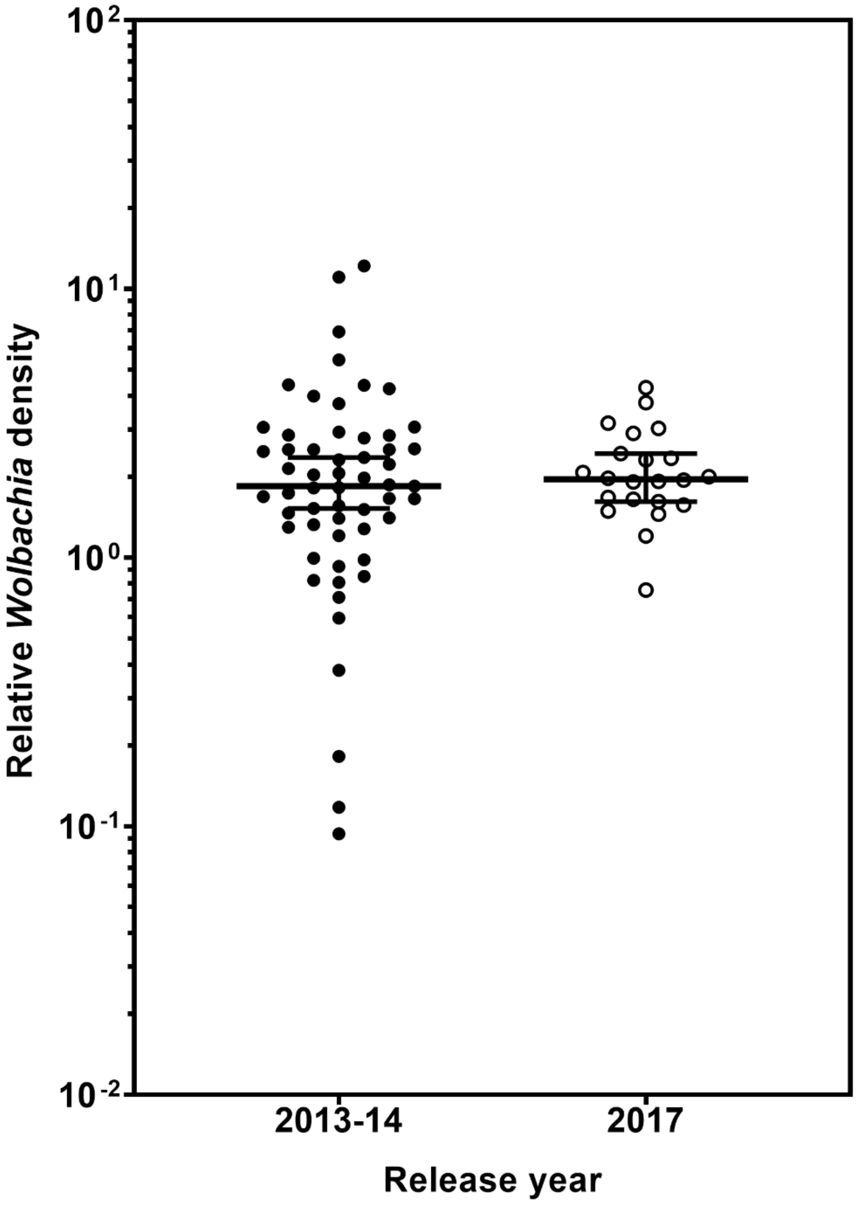

**S1 Table. Mapping statistics.**

| Sample ID | Raw reads _Lane 1 | Raw reads _Lane 2 | Read pairs after quality and adapter trimming | Number of aligned read pairs | % of mapped reads |
|---|---|---|---|---|---|
| **210290** | 34,582,326 | 34,261,303 | 48,817,765 | 43,369,702 | 88.84 |
| **210291** | 32,231,667 | 31,968,205 | 49,766,245 | 43,923,688 | 88.26 |
| **210295** | 33,580,572 | 33,244,380 | 46,137,538 | 41,094,705 | 89.07 |
| **210297** | 31,131,370 | 30,809,360 | 44,690,792 | 39,497,722 | 88.38 |
| **210298** | 30,903,897 | 30,581,705 | 48,448,379 | 43,206,264 | 89.18 |
| **210299** | 27,455,578 | 27,181,427 | 44,280,347 | 39,449,361 | 89.09 |
| **210300** | 29,400,828 | 29,080,319 | 44,257,287 | 39,327,025 | 88.86 |
| **210301** | 29,959,142 | 29,631,518 | 43,956,040 | 39,107,689 | 88.97 |
| **210304** | 32,320,363 | 31,988,121 | 42,862,574 | 38,387,721 | 89.56 |
| **210306** | 29,532,465 | 29,238,980 | 45,484,308 | 40,617,487 | 89.3 |
| **210307** | 27,631,383 | 27,344,243 | 43,284,712 | 38,791,759 | 89.62 |
| **210308** | 29,086,571 | 28,792,458 | 46,165,028 | 41,451,579 | 89.79 |
| **210309** | 31,162,958 | 30,886,606 | 49,677,557 | 44,660,124 | 89.9 |
| **210310** | 33,538,179 | 33,263,415 | 50,971,064 | 45,583,423 | 89.43 |
| **210311** | 30,444,038 | 30,183,667 | 47,816,102 | 42,728,469 | 89.36 |
| **210312** | 33,082,679 | 32,814,398 | 52,004,903 | 46,606,794 | 89.62 |
| **210313** | 27,652,952 | 27,402,701 | 43,698,630 | 38,935,479 | 89.10 |
| **210315** | 30,844,599 | 30,545,487 | 46,584,306 | 41,571,835 | 89.24 |
| **210318** | 34,698,571 | 34,346,127 | 50,984,466 | 45,600,506 | 89.44 |
| **210319** | 28,562,993 | 28,300,042 | 45,036,623 | 40,073,587 | 88.98 |
| **210320** | 33,586,176 | 33,226,843 | 45,484,408 | 40,858,644 | 89.83 |
| **210323** | 29,810,832 | 29,520,295 | 52,284,923 | 47,563,594 | 90.97 |
| **210325** | 34,343,425 | 34,074,248 | 46,562,372 | 41,831,635 | 89.84 |
| **210326** | 29,465,230 | 29,212,789 | 48,737,908 | 44,195,535 | 90.68 |
| **210327** | 31,823,328 | 31,582,004 | 49,253,762 | 44,333,311 | 90.01 |
| **210329** | 28,965,050 | 28,639,100 | 42,481,903 | 38,068,033 | 89.61 |
| **210330** | 30,294,268 | 29,888,141 | 46,080,368 | 41,043,784 | 89.07 |
| **210331** | 28,803,611 | 28,597,283 | 45,709,413 | 40,311,131 | 88.19 |
| **210332** | 30,430,171 | 30,053,350 | 48,847,663 | 43,874,971 | 89.82 |
| **210333** | 35,172,080 | 34,778,341 | 58,624,619 | 52,685,945 | 89.87 |
| **210335** | 32,188,560 | 31,805,573 | 43,615,162 | 39,044,293 | 89.52 |
| **210336** | 33,499,898 | 33,145,782 | 48,487,016 | 43,536,492 | 89.79 |
| **210338** | 29,651,131 | 29,191,005 | 43,757,822 | 38,979,468 | 89.08 |
| **Total** | 1,025,836,891 | 1,015,579,216 | 1,554,852,005 | 1,390,311,756 | 89.42 |
| **Final total** | 2,041,416,107 | | | | |

**S2 Table.** KEGG pathway enrichment analysis of 357 commonly upregulated DEGs across all time points.

| #Term ID | Term description | Observed gene count | Background gene count | False discovery rate | Matching proteins in your network (IDs) | Matching proteins in your network (labels) |
|---|---|---|---|---|---|---|
| aag00260 | Glycine, serine and threonine metabolism | 6 | 27 | 8.61E-05 | 7159.AAEL000213-PA, 7159.AAEL005336-PA, 7159.AAEL010276-PA, 7159.AAEL012578-PA, 7159.AAEL012764-PA, 7159.AAEL014426-PA | AAEL000213, AAEL005336, AAEL010276, AAEL012578, AAEL012764, AAEL014426 |
| aag01230 | Biosynthesis of amino acids | 7 | 48 | 8.61E-05 | 7159.AAEL002675-PA, 7159.AAEL003345-PA, 7159.AAEL004701-PA, 7159.AAEL005336-PA, 7159.AAEL006834-PA, 7159.AAEL008963-PA, 7159.AAEL012578-PA | AAEL002675, AAEL003345, AAEL004701, AAEL005336, AAEL006834, AAEL008963, AAEL012578 |
| aag01100 | Metabolic pathways | 22 | 716 | 0.00088 | 7159.AAEL000024-PA, 7159.AAEL000213-PA, 7159.AAEL000642-PA, 7159.AAEL001194-PA, 7159.AAEL001757-PA, 7159.AAEL002194-PA, 7159.AAEL002269-PA, 7159.AAEL002675-PA, 7159.AAEL002683-PA, 7159.AAEL003345-PA, 7159.AAEL004701-PA, 7159.AAEL005336-PA, 7159.AAEL005769-PA, 7159.AAEL005790-PA, 7159.AAEL006834-PA, 7159.AAEL008963-PA, | AAEL000024, AAEL000213, AAEL000642, AAEL001194, AAEL001757, UO, AAEL002269, AAEL002675, AAEL002683, AAEL003345, AAEL004701, AAEL005336, AAEL005769, AAEL005790, AAEL006834, AAEL008963, AAEL009679, AAEL010276, AAEL012578, AAEL013637, AAEL014426, AAEL014556 |

| | | | | | | |
|---|---|---|---|---|---|---|
| | | | | | 7159.AAEL009679-PA, 7159.AAEL010276-PA, 7159.AAEL012578-PA, 7159.AAEL013637-PA, 7159.AAEL014426-PA, 7159.AAEL015053-PB | |
| aag00220 | Arginine biosynthesis | 3 | 13 | 0.008 | 7159.AAEL002675-PA, 7159.AAEL003345-PA, 7159.AAEL004701-PA | AAEL002675, AAEL003345, AAEL004701 |
| aag00232 | Caffeine metabolism | 2 | 3 | 0.0107 | 7159.AAEL002194-PA, 7159.AAEL002683-PA | UO, AAEL002683 |
| aag00350 | Tyrosine metabolism | 3 | 17 | 0.0107 | 7159.AAEL000024-PA, 7159.AAEL008963-PA, 7159.AAEL013637-PA | AAEL000024, AAEL008963, AAEL013637 |
| aag01200 | Carbon metabolism | 5 | 78 | 0.0172 | 7159.AAEL005336-PA, 7159.AAEL005790-PA, 7159.AAEL010276-PA, 7159.AAEL012578-PA, 7159.AAEL014426-PA | AAEL005336, AAEL005790, AAEL010276, AAEL012578, AAEL014426 |
| aag00330 | Arginine and proline metabolism | 3 | 26 | 0.0234 | 7159.AAEL000213-PA, 7159.AAEL002675-PA, 7159.AAEL006834-PA | AAEL000213, AAEL002675, AAEL006834 |
| aag00330 | Arginine and proline metabolism | 3 | 26 | 0.0301 | 7159.AAEL003116-PA, 7159.AAEL005575-PA, 7159.AAEL008108-PA | AAEL003116, AAEL005575, AAEL008108 |
| aag04745 | Phototransduction - fly | 3 | 30 | 8.61E-05 | 7159.AAEL000213-PA, 7159.AAEL005336-PA, 7159.AAEL010276-PA, 7159.AAEL012578-PA, 7159.AAEL012764-PA, 7159.AAEL014426-PA | AAEL000213, AAEL005336, AAEL010276, AAEL012578, AAEL012764, AAEL014426 |

**S3 Table. Unmapped downregulated DEGs which were non-coding RNA from *Aedes aegypti* with different release years.**

| *Aedes aegypti* release year | | |
|---|---|---|
| **Aae.wMel$_{2011}$** | **Aae.wMel$_{2013/2014}$** | **Aae.wMel$_{2017}$** |
| LOC110674171, LOC110677214, LOC110674245, LOC110676610, LOC110674560, LOC110676459, LOC110678327, LOC110679860, LOC110678696, LOC110676194, LOC110679144, LOC110675307, LOC110677893, LOC110676333, LOC110674112 and LOC110674372 | LOC110679264, LOC110675798, LOC110679860, LOC110675419, LOC110678052, LOC110679845, LOC110678327, LOC110681560, LOC110677252, LOC110679144 and LOC110674112 | LOC110674426, LOC110675307, LOC110679848, LOC110677249, LOC110675921, LOC110679654, LOC110678978, LOC110679077, LOC110676416, LOC110675448, LOC110676459, LOC110680245, LOC110675798, LOC110674875, LOC110675739, LOC110676610, LOC110676947, LOC110674156, LOC110676068, LOC110674372, LOC110675480, LOC110680326, LOC110679406, LOC110678052, LOC110680549, LOC110678657, LOC110679051, LOC110678740, LOC110677977, LOC110675419, LOC110679845, LOC110676013, LOC110679264 and LOC110676333 |

**S4 Table. Unmapped downregulated DEGs in Aae.wMel$_{2011}$ mosquitoes which are not ncRNA.**

| Entrez Gene ID | Description/ Name | Function | Reference |
|---|---|---|---|
| 110678281 | pfam: Dimer_Tnp_hAT; hAT family C-terminal dimerisation region | Protein bares much similarity to zinc finger or transposase proteins | http://pfam.xfam.org/family/Dimer_Tnp_hAT |
| 110676828 | replication factor C subunit 3-like / Rfc5p, putative | DNA-directed DNA polymerase | https://www.ncbi.nlm.nih.gov/gene/?term=110676828 |
| 110674591 | SMC_N; RecF/RecN/SMC N terminal domain | Structural maintenance of chromosomes_N Superfamily | https://www.ncbi.nlm.nih.gov/gene/?term=110674591 |
| 110680308 | Protein Optix-like | May be involved in head or eye development; development of the clypeolabrum and several head sensory organs | https://www.uniprot.org/uniprot/Q95RW8 |
| 110675146 | DNA Binding<br>Homologous to Deformed epidermal autoregulatory factor 1-like | When secreted, behaves as an inhibitor of cell proliferation, by arresting cells in the G0 or G1 phase. | https://www.uniprot.org/uniprot/A0A6I8U8T1<br>https://www.uniprot.org/uniprot/O75398 |
| 110676629 | Uncharacterised | | https://www.ncbi.nlm.nih.gov/gene/?term=110676629 |
| 110679076 | Uncharacterised | | https://www.ncbi.nlm.nih.gov/gene/?term=110679076 |
| 110680643 | Carcinine transporter-like | CarT expression in photoreceptors is necessary and sufficient for fly vision and behavior. | https://www.ncbi.nlm.nih.gov/pmc/articles/PMC4739767/ |
| 110679455 | Protein commissureless 2 homolog | Essential for nerve cord development. Functions downstream of fra to control axon guidance across the central nervous system (CNS) midline | https://www.ncbi.nlm.nih.gov/gene/?term=110679455,<br>https://www.uniprot.org/uniprot/Q9VUT8 |
| CFI06_mgr02 | 16S ribosomal RNA | Translation process | https://www.ncbi.nlm.nih.gov/gene/?term=CFI06_mgr02 |
| 110679193 | Putative oxidoreductase GLYR1 homolog | Oxidoreductase activity, positive regulation of histone acetylation, positive regulation of transcription by RNA polymerase II | https://www.ncbi.nlm.nih.gov/gene/?term=110679193,<br>https://www.uniprot.org/uniprot/Q8T079 |
| 110678839 | Zinc finger C4H2 domain-containing protein-like | Plays a role in interneurons differentiation | https://www.ncbi.nlm.nih.gov/gene/?term=110678839,<br>https://www.uniprot.org/uniprot/Q9NQZ6 |
| 110675237 | Apolipoprotein D-like | Apolipoprotein D (ApoD) is an extracellular glycoprotein of the lipocalin protein family, involved in different functions such as immune response, cell proliferation | https://www.ncbi.nlm.nih.gov/gene/?term=110675237,<br>https://www.intechopen.com/books/advances-in-lipoprotein-research/apolipoprotein-d |

| | | | |
|---|---|---|---|
| | | regulation, chemoreception, retinoid metabolism, axon growth, and proteolysis regulation. | |
| 110675476 | Uncharacterised | | https://www.ncbi.nlm.nih.gov/gene/?term=110675476 |
| 110680748 | Zinc finger protein 569-like | ZNF569 protein may act as a transcriptional repressor that suppresses MAPK signaling pathway to mediate cellular functions. | https://www.ncbi.nlm.nih.gov/gene/?term=110680748, https://pubmed.ncbi.nlm.nih.gov/16793018/ |
| 110676559 | BEN domain | BEN domain mediates protein-DNA and protein-protein interactions during chromatin organisation and transcription. The presence of BEN domains in a poxviral early virosomal protein and in polydnaviral proteins also suggests a possible role for them in organization of viral DNA during replication or transcription. | https://www.ncbi.nlm.nih.gov/gene/?term=110676559, https://www.ncbi.nlm.nih.gov/pmc/articles/PMC2477736/ |
| 110676930 | Testis-specific zinc finger protein topi-like | The Drosophila aly-class meiotic arrest loci are essential for activation of transcription of many differentiation-specific genes, as well as several genes important for meiotic cell cycle progression, thus linking meiotic cell cycle progression to cellular differentiation during spermatogenesis. | https://www.ncbi.nlm.nih.gov/gene/?term=110676930, https://pubmed.ncbi.nlm.nih.gov/15084455/ |
| 110680210 | Bromodomain adjacent to zinc finger domain protein 1A-like | Chromatin remodeling, DNA-dependent DNA replication, histone acetylation and regulation of transcription by RNA polymerase II | https://www.ncbi.nlm.nih.gov/gene/?term=110680210, |
| 110679465 | Uncharaterised | | https://www.ncbi.nlm.nih.gov/gene/?term=110679465 |
| 110680634 | PAB-dependent poly(A)-specific ribonuclease subunit PAN3-like | deadenylation-dependent decapping of nuclear-transcribed mRNA Source: GO_Central<br>mRNA processing, nuclear-transcribed mRNA poly(A) tail shortening, positive regulation of cytoplasmic mRNA processing body assembly | https://www.ncbi.nlm.nih.gov/gene/?term=110680634, https://www.uniprot.org/uniprot/B7Q0Q0 |

**S5 Table. Unmapped downregulated DEGs from Aae.wMel$_{2013/2014}$ mosquitoes.**

| Entrez Gene ID | Description | Function | Reference |
|---|---|---|---|
| LOC110678585 | Paired box protein Pax-6-like | Involved in eye morphogenesis | https://www.ncbi.nlm.nih.gov/gene/?term=110678585, https://www.uniprot.org/uniprot/O18381 |
| LOC110674017 | Polyprenol reductase-like | N-Glycan biosynthesis | https://www.ncbi.nlm.nih.gov/gene/?term=110674017 |
| LOC110676930 | Testis-specific zinc finger protein topi-like | The Drosophila aly-class meiotic arrest loci are essential for activation of transcription of many differentiation-specific genes, as well as several genes important for meiotic cell cycle progression, thus linking meiotic cell cycle progression to cellular differentiation during spermatogenesis. | https://www.ncbi.nlm.nih.gov/gene/?term=110676930, https://pubmed.ncbi.nlm.nih.gov/15084455/ |
| LOC110675182 | Uncharcterised, BEN; BEN domain | ?Organization of viral DNA during replication or transcription | https://www.ncbi.nlm.nih.gov/gene/?term=110675182, https://www.ncbi.nlm.nih.gov/pmc/articles/PMC2477736/ |
| LOC110678581 | Uncharacterized, MADF_DNA_bdg; Alcohol dehydrogenase transcription factor Myb/SANT-like | Transcription factor | https://www.ncbi.nlm.nih.gov/gene/?term=110678581, https://www.genome.jp/kegg-bin/get_htext |
| LOC110676965 | Zinc finger BED domain-containing protein 1-like | Protein dimerization activity | https://www.ncbi.nlm.nih.gov/gene/?term=110676965, https://www.uniprot.org/uniprot/F6QDB5 |
| LOC110674313 | Zinc finger protein 845-like | Transcription factor | https://www.ncbi.nlm.nih.gov/gene/?term=110674313, https://www.genome.jp/kegg-bin/get_htext |
| LOC110678090 | ATP-dependent DNA helicase PIF1-like | DNA replication proteins | https://www.ncbi.nlm.nih.gov/gene/?term=110678090, https://www.genome.jp/kegg-bin/get_htext?aag00001+5580192 |
| CFI06_mgr02 | 16S ribosomal RNA | Translation process | https://www.ncbi.nlm.nih.gov/gene/?term=CFI06_mgr02 |
| LOC110676559 | BEN domain | BEN domain mediates protein-DNA and protein-protein interactions during chromatin organisation and transcription. The presence of BEN domains in a poxviral early virosomal protein and in polydnaviral proteins also suggests a possible role for them in organization of viral DNA during replication or transcription. | https://www.ncbi.nlm.nih.gov/gene/?term=110676559, https://www.ncbi.nlm.nih.gov/pmc/articles/PMC2477736/ |
| LOC110675146 | Deformed epidermal autoregulatory factor 1-like | Transcription factor | https://www.uniprot.org/uniprot/O75398 |
| LOC110676076 | Mantle protein-like | Protein Blast identified 98% similarity to Vajk2, which playes a role in chitin-based cuticle development in *Drosophila melanogaster.* | https://www.ncbi.nlm.nih.gov/gene/?term=110676076, https://www.uniprot.org/uniprot/Q8SZM2 |

**S1 Box. Significantly enriched GO terms pertaining to biological processes and molecular functions in Aae.wMel$_{2017}$ mosquitoes.**

**A) Biological processes**

GO:0006270 DNA replication initiation
GO:0006260 DNA replication
GO:0060070 canonical Wnt signaling pathway
GO:0000724 double-strand break repair via homologous recombination
GO:0006281 DNA repair
GO:0006979 response to oxidative stress
GO:0006334 nucleosome assembly
GO:0007275 multicellular organism development
GO:0042438 melanin biosynthetic process
GO:0060179 male mating behavior
GO:0071139 resolution of recombination intermediates
GO:0007018 microtubule-based movement

**B) Molecular functions**

GO:0005524 ATP binding
GO:0004672 protein kinase activity
GO:0003684 damaged DNA binding
GO:0004601 peroxidase activity
GO:0003678 DNA helicase activity
GO:0042813 Wnt-activated receptor activity
GO:0004674 protein serine/threonine kinase activity
GO:0008270 zinc ion binding
GO:0003777 microtubule motor activity

**S6 Table. Unmapped downregulated DEGs in Aae.wMel$_{2017}$ mosquitoes.**

| Gene ID | Description | Function | Reference |
|---|---|---|---|
| LOC110675237 | 40S ribosomal protein S17 | Translation | https://www.ncbi.nlm.nih.gov/gene/?term=110680939, https://www.uniprot.org/uniprot/Q52UT2 |
| LOC110678090 | apolipoprotein D-like | | https://www.ncbi.nlm.nih.gov/gene/?term=110675237 |
| LOC110675182 | ATP-dependent DNA helicase PIF1-like | DNA-dependent ATPase and 5'-3' DNA helicase required for the maintenance of both mitochondrial and nuclear genome stability | https://www.ncbi.nlm.nih.gov/gene/?term=110678090, https://www.uniprot.org/uniprot/Q9H611 |
| LOC110677148 | BEN domain | | https://www.ncbi.nlm.nih.gov/gene/?term=110675182%5Buid%5D |
| LOC110680907 | cell division cycle protein 20 homolog | This protein is involved in the pathway protein ubiquitination, which is part of Protein modification. | https://www.ncbi.nlm.nih.gov/gene/?term=110677148, https://www.uniprot.org/uniprot/Q12834 |
| LOC110674064 | CREB-regulated transcription coactivator 1-like | cAMP-responsive element binding protein (CREB) has been well known as one of the best studied inducible eukaryotic transcription factors. | https://www.ncbi.nlm.nih.gov/gene/?term=110680907, https://pubmed.ncbi.nlm.nih.gov/25896045/ |
| LOC110680411 | DNA repair protein XRCC3-like | Involved in the homologous recombination repair (HRR) pathway of double-stranded DNA, thought to repair chromosomal fragmentation, translocations, and deletions. | https://www.ncbi.nlm.nih.gov/gene/?term=110674064, https://www.uniprot.org/uniprot/O43542 |
| LOC110674134 | Domain of unknown function (DUF4806) | | https://www.ncbi.nlm.nih.gov/gene/?term=110680411 |
| LOC110680252 | dynein heavy chain 10, axonemal | Force generating protein of respiratory cilia. Produces force towards the minus ends of microtubules. Dynein has ATPase activity | https://www.ncbi.nlm.nih.gov/gene/?term=110674134, https://www.uniprot.org/uniprot/Q8IVF4 |
| LOC110674232 | E3 SUMO-protein ligase PIAS2-like | We show that the mosquito SUMOylation pathway plays a broadly antiviral role against a wide range of clinically important arboviruses, including Zika, Semliki Forest, and Bunyamwera viruses. SUMO important in post-translational modification. | https://www.ncbi.nlm.nih.gov/gene/?term=110680252, https://journals.plos.org/plospathogens/article?id=10.1371/journal.ppat.1009134, |
| LOC110676122 | Ephexin Pleckstrin homology (PH) domain | PH domains play a role in recruiting proteins to different membranes, thus targeting them to appropriate cellular compartments or enabling them to interact with other components of the signal transduction pathways. | https://www.ncbi.nlm.nih.gov/gene/?term=110674232, https://www.ebi.ac.uk/interpro/entry/InterPro/IPR001849/ |
| LOC110673995 | fer-1-like protein 6 | transcription factors as being involved in *Odorant receptor* regulation | https://www.ncbi.nlm.nih.gov/gene/?term=110676122, https://bmcgenomics.biomedcentral.com/articles/10.1186/s12864-020-07336-w |

| | | | |
|---|---|---|---|
| LOC110679476 | FLYWCH zinc finger domain | This domain was first characterised in Drosophila Modifier of mdg4 proteins, Mod(mgd4), putative chromatin modulators involved in higher order chromatin domains. | https://www.ebi.ac.uk/interpro/entry/InterPro/IPR007588/ |
| LOC110678281 | hAT family C-terminal dimerisation region | the protein bares much similarity to zinc finger or transposase proteins. | https://www.ncbi.nlm.nih.gov/gene/?term=110679476, http://pfam.xfam.org/family/Dimer_Tnp_hAT |
| LOC110674011 | hAT family C-terminal dimerisation region | | https://www.ncbi.nlm.nih.gov/gene/?term=110678281 |
| LOC110675487 | Headcase protein-like | Required for imaginal cell differentiation, may be involved in hormonal responsiveness during metamorphosis (34), headcase was identified in a screen as a regulator of the siRNA pathway (35) | https://www.ncbi.nlm.nih.gov/gene/?term=110674011 |
| LOC110676735 | histone deacetylase Rpd3-like | Histone deacetylation plays an important role in transcriptional regulation, cell cycle progression, DNA damage response, osmotic stress response and developmental events, | https://www.ncbi.nlm.nih.gov/gene/?term=110675487, https://www.uniprot.org/uniprot/P32561 |
| LOC110675977 | histone H4 | Sense and antisense histone 4-derived piRNAs accumulate in an Ago3/Piwi5 ping-pong-dependent fashion | https://www.ncbi.nlm.nih.gov/gene/?term=110676735, https://www.jimmunol.org/content/jimmunol/190/2/650.full.pdf?with-ds=yes |
| LOC110680300 | lactosylceramide 1,3-N-acetyl-beta-D-glucosaminyltransferase | sphingolipid biosynthesis, plays a key role in the synthesis of lacto- or neolacto-series carbohydrate chains on glycolipids | https://www.ncbi.nlm.nih.gov/gene/?term=110675977, https://journals.plos.org/plosone/article/file?type=supplementary&id=info:doi/10.1371/journal.pone.0155616.s005 Massive Shift in Gene Expression during Transitions between Developmental Stages of the Gall Midge, Mayetiola Destructor |
| LOC110678193 | leucine-rich repeat [structural motif] | | https://www.ncbi.nlm.nih.gov/gene/?term=110680300 |
| LOC110679126 | lipid storage droplets surface-binding protein 2-like | Several RNA viruses use host LDs at different steps of their life cycle | https://www.ncbi.nlm.nih.gov/gene/?term=110678193, https://www.ncbi.nlm.nih.gov/pmc/articles/PMC3268388/ |
| LOC110678581 | LRR_AMN1; leucine-rich repeat [structural motif] | Immune | https://www.ncbi.nlm.nih.gov/gene/?term=110679126 |
| LOC110677913 | MADF_DNA_bdg; Alcohol dehydrogenase | regulation of transcription, DNA-templated | https://www.ncbi.nlm.nih.gov/gene/?term=110678581, https://www.uniprot.org/uniprot/Q9LV59 |

| | | | |
|---|---|---|---|
| | transcription factor Myb/SANT-like | | |
| LOC110679883 | Major facilitator superfamily domain-containing protein 10 | apoptotic process, sodium-independent organic anion transport | https://www.ncbi.nlm.nih.gov/gene/?term=110677913, https://www.uniprot.org/uniprot/Q14728 |
| LOC110679883 | Mucin-5AC-like | Amino sugar and nucleotide sugar metabolism | https://www.ncbi.nlm.nih.gov/gene/?term=110679883, https://www.genome.jp/kegg-bin/get_htext |
| LOC110680249 | oocyte zinc finger protein XlCOF22-like | Transcription, Transcription regulation | https://www.ncbi.nlm.nih.gov/gene/?term=110680249 |
| LOC110680634 | PAB-dependent poly(A)-specific ribonuclease subunit PAN3-like | mRNA surveillance and transport factors | https://www.ncbi.nlm.nih.gov/gene/?term=110680634, https://www.genome.jp/kegg-bin/get_htext?aag03019+5577183 |
| LOC110678585 | Paired box protein Pax-6-like | Involved in eye morphogenesis and adult development | https://www.ncbi.nlm.nih.gov/gene/?term=110678585, https://www.uniprot.org/uniprot/O18381 |
| LOC110676083 | Probable E3 ubiquitin protein ligase DRIPH | This protein is involved in the pathway protein ubiquitination, which is part of Protein modification. | https://www.ncbi.nlm.nih.gov/gene/?term=110676083, https://www.uniprot.org/uniprot/Q9LS86 |
| LOC110680850 | Proline-rich protein 36 | This gene encodes a large protein of unknown function that contains internal regions of low complexity sequence. Alternative splicing results in multiple transcript variants. The transcript structure of the protein-coding variant at this locus is conserved between human and mouse | https://www.ncbi.nlm.nih.gov/gene/?term=110680850, https://www.genecards.org/cgi-bin/carddisp.pl?gene=PRR36 |
| LOC110679455 | Protein commissureless 2 homolog | Essential for nerve cord development. Functions downstream of fra to control axon guidance across the central nervous system (CNS) midline | https://www.ncbi.nlm.nih.gov/gene/?term=110679455, https://www.uniprot.org/uniprot/Q9VUT8 |
| LOC110676568 | Putative defense protein Hdd11-like | May have antimicrobial activity, defense response to bacterium, defense response to protozoan, innate immune response | https://www.ncbi.nlm.nih.gov/gene/?term=110676568, https://www.uniprot.org/uniprot/Q86RS3 |
| LOC110680234 | RecF/RecN/SMC N terminal domain | function together with other proteins in a range of chromosomal transactions, including chromosome condensation, sister-chromatid cohesion, recombination, DNA repair and epigenetic silencing of gene expression | https://www.ncbi.nlm.nih.gov/gene/?term=110680234, https://www.ebi.ac.uk/interpro/entry/InterPro/IPR003395/ |
| LOC110676828 | replication factor C subunit 3-like | May be involved in DNA replication and thus regulate cell proliferation | https://www.ncbi.nlm.nih.gov/gene/?term=110676828, https://www.uniprot.org/uniprot/Q8VXX4 |
| LOC110674773 | Sec63; Sec63 Brl domain | Proteins destined for the secretory pathway are initially translocated across the membrane of the endoplasmic | https://www.ncbi.nlm.nih.gov/gene/?term=110674773 |

| | | | |
|---|---|---|---|
| | | reticulum (ER)4 at pore-forming structures known as translocons. A substantial body of data indicates that Sec63p and Kar2p contribute directly to the driving force for post-translational translocation, with the ATPase activity of Kar2p being activated by the luminal J-domain within Sec63p (9, 10). basic function in translocation | |
| LOC110678733 | serine proteinase stubble | Trypsin | https://www.ncbi.nlm.nih.gov/gene/?term=110678733, https://www.genome.jp/dbget-bin/www_bget?aag:5572015 |
| LOC110680822 | set1/Ash2 histone methyltransferase complex subunit ASH2-like | Transcriptional regulator | https://www.ncbi.nlm.nih.gov/gene/?term=110680822, https://www.uniprot.org/uniprot/Q9UBL3 |
| LOC110677032 | short stature homeobox protein 2-like | May be a growth regulator and have a role in specifying neural systems involved in processing somatosensory information, as well as in face and body structure formation. | https://www.ncbi.nlm.nih.gov/gene/?term=110677032, https://www.uniprot.org/uniprot/O60902 |
| LOC110679408 | Tc5 transposase DNA-binding domain | DNA-binding | https://www.ncbi.nlm.nih.gov/gene/?term=110679408, https://www.uniprot.org/uniprot/Q23E04 |
| LOC110675561 | tektin-1 | Tektins are insoluble α-helical proteins essential for the construction of cilia and flagella and are found throughout the eukaryotes apart from higher plants. | https://www.ncbi.nlm.nih.gov/gene/?term=110675561, https://genomebiology.biomedcentral.com/articles/10.1186/gb-2008-9-7-229 |
| LOC110676106 | tubulin alpha-8 chain-like | Tubulin is the major constituent of microtubules. | https://www.ncbi.nlm.nih.gov/gene/?term=110676106, https://www.uniprot.org/uniprot/Q9NY65 |
| LOC110679470 | Tudor domain | TUDOR-domain containing (Tudor) proteins facilitate piRNA biogenesis in Drosophila melanogaster and other model organisms. | https://www.ncbi.nlm.nih.gov/gene/?term=110679470, https://www.researchgate.net/publication/329844811_The_Tudor_protein_Veneno_assembles_the_ping-pong_amplification_complex_that_produces_viral_piRNAs_in_Aedes_mosquitoes |
| LOC110681183 | WASH complex subunit 2-like | protein localization to endosome, protein transport, retrograde transport, endosome to Golgi | https://www.ncbi.nlm.nih.gov/gene/?term=110681183, https://www.uniprot.org/uniprot/Q6PGL7 |
| LOC110674056 | Werner syndrome ATP-dependent helicase-like | May play an important role in the dissociation of joint DNA molecules that can arise as products of homologous recombination, at stalled replication forks or during DNA repair. Alleviates stalling of DNA polymerases at the site of DNA lesions. Important for genomic integrity. Plays a role in the formation of DNA replication focal centers; | https://www.ncbi.nlm.nih.gov/gene/?term=110674056, https://www.uniprot.org/uniprot/Q14191 |

| | | | |
|---|---|---|---|
| | | stably associates with foci elements generating binding sites for RP-A (By similarity). Plays a role in double-strand break repair after gamma-irradiation. | |
| LOC110680251 | XPG domain containing | DNA repair | https://www.ncbi.nlm.nih.gov/gene/?term=110680251, https://www.uniprot.org/uniprot/A0A1S4EVT1 |
| LOC110676965 | zinc finger BED domain-containing protein 1-like | DNA binding | https://www.ncbi.nlm.nih.gov/gene/?term=110676965, https://www.uniprot.org/uniprot/F6QDB5 |
| LOC110681552 | zinc finger protein 148-like | Involved in transcriptional regulation. Represses the transcription of a number of genes including gastrin, stromelysin and enolase. Binds to the G-rich box in the enhancer region of these genes. | https://www.ncbi.nlm.nih.gov/gene/?term=110681552, https://www.uniprot.org/uniprot/Q9UQR1 |
| LOC110680748 | zinc finger protein 569-like | May be involved in transcriptional regulation | https://www.ncbi.nlm.nih.gov/gene/?term=110680748, https://www.uniprot.org/uniprot/Q5MCW4#function |
| LOC110674313 | zinc finger protein 845-like | May be involved in transcriptional regulation. | https://www.ncbi.nlm.nih.gov/gene/?term=110674313, https://www.genecards.org/cgi-bin/carddisp.pl?gene=ZNF845 |
| LOC110679831 | zinc finger protein DZIP1L | Involved in primary cilium formation | https://www.ncbi.nlm.nih.gov/gene/?term=110679831, https://www.uniprot.org/uniprot/Q8IYY4 |
